# A versatile method to pattern surfaces within microfluidic devices

**DOI:** 10.64898/2026.02.19.706879

**Authors:** Kate Collins, Claire E. Stanley, Thomas E. Ouldridge

## Abstract

Microfluidic devices with surface-bound biomolecular patterns enable localised detection arrays, enzymatic catalysis, and gene expression. Photolithography is a well-established method to pattern open surfaces with high spatial control. However, patterning enclosed microfluidic channels remains technically challenging. Such capability would enable *in situ* surface modification and precise pattern alignment to channel geometries. Here, we present a photolithographic method using commercially available reagents to pattern sealed microfluidic devices. We first coat surfaces with (3-Aminopropyl)triethoxysilane (APTES) to bond microfluidic chips and provide surface amine groups onto which photocleavable polyethylene glycol (PC PEG) compounds are bound. UV exposure using standard photolithography equipment selectively deprotects the amine groups, which can subsequently bind amine-reactive cargos. We demonstrate the versatility of this method by patterning both poly(dimethylsiloxane) (PDMS) and glass surfaces with diverse cargoes: DNA, proteins, and gold nanoparticles. We also compare covalent versus noncovalent DNA patterning. Covalently bound DNA patterns are denser and could be used for sequence-specific target DNA capture. However, noncovalently bound DNA yielded higher cell-free gene expression from surface-bound GFP templates.

## Introduction

Since their invention in the 1990s, microfluidic devices have evolved from simple channels to complex miniaturized experimental platforms with applications in biology, biotechnology, and chemistry.^1^ Microfluidic devices are typically formed by bonding a structured poly(dimethylsiloxane) (PDMS) top layer containing micrometer-sized features to a flat substrate such as glass. PDMS is an optically transparent material, so enclosed systems can be monitored by microscopy – for example, to observe cellular behaviour or chemical reaction networks in real time.^2,3^ On the microscale, fluid flow is predominantly laminar,^4^ allowing more precise fluid transport, mixing and encapsulation than traditional benchtop platforms.^5^ Miniaturization has another consequence: as reactors are scaled down, the surface area-to-volume ratio increases, inflating the effects of surface interactions. Therefore, engineering surfaces within microfluidic devices is a key consideration in device design. ^6,7^ Microfluidic surfaces have been chemically modified to alter surface hydrophobicity,^8,9^ reduce nonspecific adsorption,^10,11^ and anchor cells^12–14^ or biomolecules such as antibodies ^15,16^ and DNA.^17^

Spatially localizing such surface modification provides additional avenues to engineer sophisticated microfluidic platforms. For instance, protein arrays have been used for onchip immunoassays,^18,19^ cell isolation,^16,20^ and localized enzymatic reaction cascades.^21–23^ Surface-bound cellular patterns have been used to probe cellular interactions and environmental responses.^20,24^ Similarly, DNA patterns have been used for microfluidic DNA sequence detection platforms^25^ and cell-free gene expression chambers.^26,27^

Patterning surfaces within microfluidic devices is challenging. Two approaches have emerged: patterning open surfaces before bonding devices, while surfaces are accessible, or using non-contact methods to modify surfaces after devices are bonded.^28^ In the first case, open surfaces can be patterned using well-established methods such as microcontact printing^18,29^ and droplet printing.^10,30^ While the patterning step is straightforward, the modified surfaces are often incompatible with plasma-based bonding, and require bespoke device assembly methods such as scaffolds^17,31^ or lamination.^32^ In addition, patterns cannot be precisely aligned to microfluidic features during the bonding step. Conversely, methods based on post-bonding modifications bypass bonding and alignment issues, at the cost of surface access. Patterning can be achieved by co-flowing liquid streamlines containing surface-binding reagents^12,33,34^ or physically blocking areas using valves.^10,30^ These approaches are however limited in pattern shape and complexity, and require complex microfluidic devices.

Surfaces can also be patterned by photolithography, where light is projected onto photoreactive substrates.^35^ Photolithography can be performed within sealed devices,^24^ does not require specific microfluidic features such as valves, and can print arbitrary shapes. ^17,31^ Projected light can also be aligned to microchannel features, for example by using digital micromirror devices (DMD) integrated into microscopes.^36^ Microfluidic surfaces have notably been patterned using photo-reactive poly(ethylene glycol) (PEG) diacrylate,^11^ thiol-ene chemistry,^37^ photo-reactive DNA^17,36^ and photocleavable PEG monolayers.^31,38^ There are, however, key limitations to the broader implementation of photolithography-based microfluidic surface patterning. First, existing methods remain technically challenging as they often require bespoke chemical synthesis, photolithographic setups, and device holders.^24,31,39^ Second, current techniques exhibit limited versatility, with restrictions in material compatibility, patterned cargo, and photochemical mechanism.^11,16,23,31,40–44^

Here, we seek to develop an accessible, modular approach to pattern surfaces within microfluidic channels using photolithography. To do so, we base our approach on modifying surfaces with (3-Aminopropyl)triethoxysilane (APTES). APTES coatings have been used as adhesives to bind PDMS to glass, thermoplastics,^45–48^ polycarbonate membranes,^49^ and even metals.^50^ APTES contains an amine group, resulting in a positively charged surface. Cells,^51^ proteins^13,14,52,53^ and DNA^54,55^ have been immobilized on APTES-coated surfaces through noncovalent interactions. In addition, the amine group can be covalently modified with N-hydroxysuccinimide (NHS) esters,^56–58^ glutaraldehyde^59,60^ and NHS-(1-Ethyl-3-[3-dimethylaminopropyl]carbodiimide) (EDC) conjugation.^18,61,62^ Although APTES coatings have been used to pattern surfaces by microcontact printing,^18,29,52^ plasma micropatterning,^15,63^ and photoresist patterning,^53,64^ their use for photopatterning remains largely unexplored. One notable exception involved patterning a polymer inside APTES-coated glass capillaries to immobilize cells. ^65^

Recently, a range of amine-reactive, photocleavable (PC) PEG compounds have become commercially available, featuring NHS-esters on one end and ‘head’ groups (such as biotin or azide) on the other.^66^ In this work, we explore the use of such PC PEG compounds to pattern APTES within microfluidic devices by photolithography (Fig. 1). We investigate the versatility of this approach by patterning different materials and functionalizing both the PC PEG head groups and the UV-exposed APTES patterns. While both PC PEG and APTES patterns can be covalently functionalized with cargos, APTES patterns can also bind cargos noncovalently. We subsequently compare the functionality of covalently and noncovalently bound DNA patterns.

**Figure 1:**
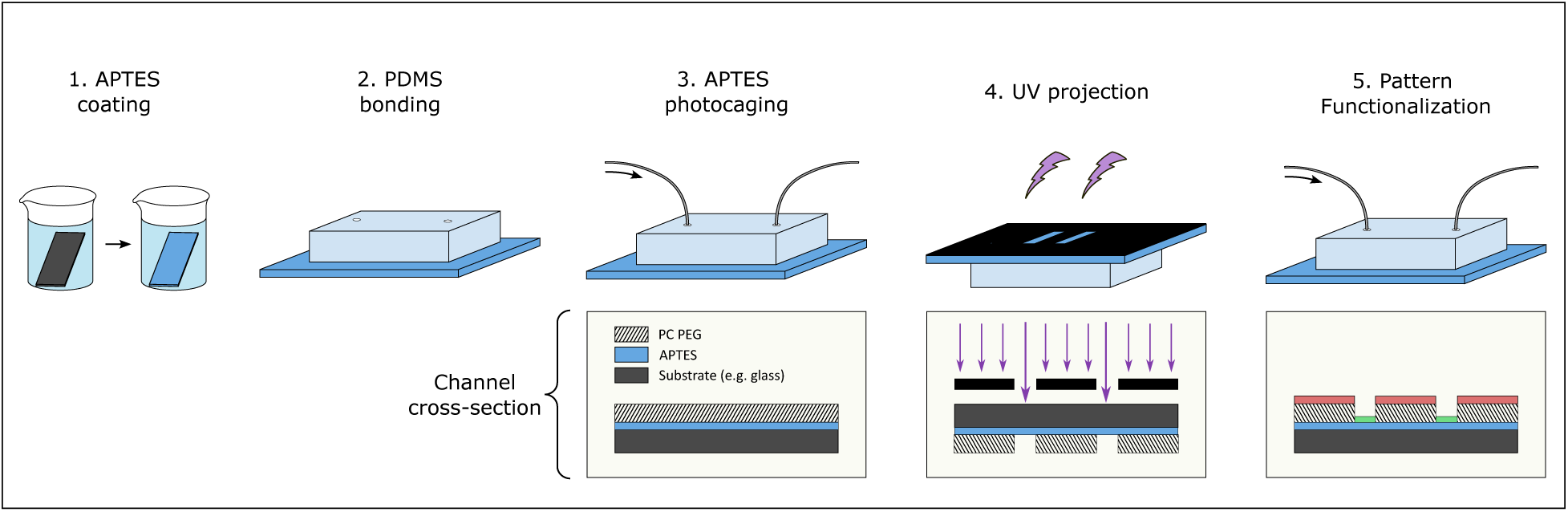
Overview of APTES-based microfluidic patterning. A substrate (such as glass) is coated with APTES, which acts as an adhesive to bind a structured PDMS top layer. The assembled device is then loaded with amine-reactive photocleavable PEG (PC PEG) to photocage APTES, and subsequently exposed to a patterned UV signal. In UV-exposed areas, PC PEG is cleaved, deprotecting the APTES amine groups. The resulting patterns can be functionalized using the PC PEG head group (negative tone patterns, red) or the APTES amine groups (positive tone patterns, green).

## Results

A prerequisite for the patterning strategy was to produce APTES coatings that both mediated device bonding and amine reactivity within sealed microchannels. First, we incubated plasma-treated glass or PDMS substates in 1% APTES, upon which a plasma-treated PDMS top layer was bound to create a series of microchannels (Fig. 2A). The resulting devices could not be manually peeled apart. In order to test if this protocol led to the formation of aminated surfaces, we first carried out an X-ray photoelectron spectroscopy (XPS) analysis of APTES-coated glass surfaces. In comparison with untreated controls, APTES-modified surfaces had detectable NH^+^ and NH_2_ peaks (Fig. 2B, Fig. S1), alongside a relative increase in carbon and nitrogen content (Table S1).

**Figure 2:**
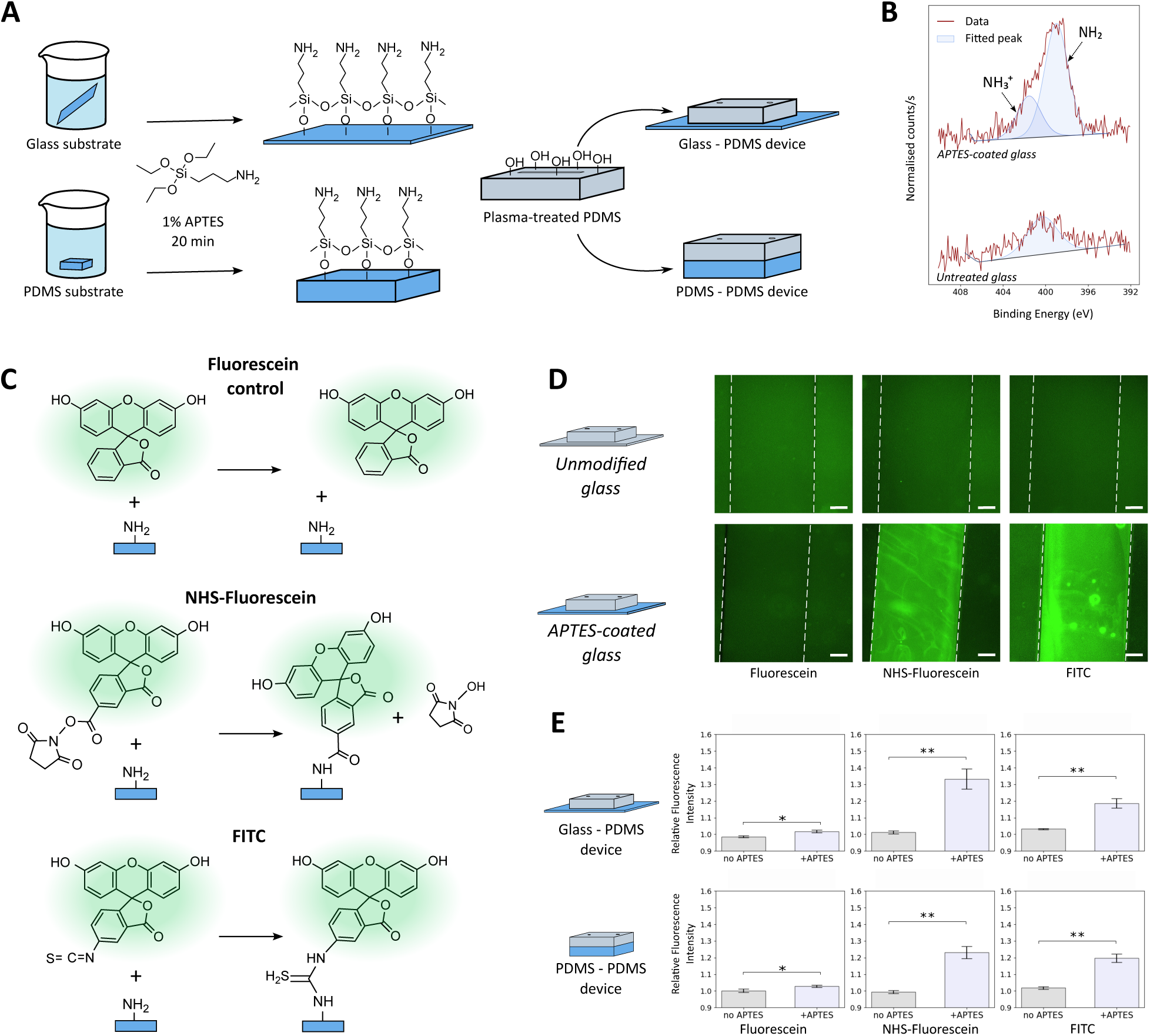
APTES surface characterization. **(A)** Glass or PDMS substrates were pre-coated with APTES, and then bound to PDMS top layers. **(B)** XPS N1s high-resolution scans. APTES coating led to surface amination. **(C)** Fluorescein derivative assay system to detect amine groups within sealed microfluidic channels by fluorescence. Fluorescein alone does not bind amine groups, while NHS-Fluorescein and FITC bind amine groups through different mechanisms. **(D)** Representative images of fluorescein assays on unmodified and APTES-coated glass substrates, showing fluorescence only in the presence of APTES and amine-reactive groups. Scale bars: 100 µm. **(E)** Fluorescein derivatives assay on glass and PDMS substrates within microfluidic devices. Amine-reactive probes adhered to microchannel surfaces, indicating the presence of APTES amine groups (Mann-Whitney U test, n=6-8 for glass-PDMS devices and 4-6 for PDMS-PDMS devices).

We then tested the presence of surface amine groups within sealed microfluidic devices, using simple straight-channel geometries (Fig. S2). We designed an assay based on fluorescein derivatives, where NHS-Fluorescein and fluorescein isothiocyanate (FITC) react with amine groups through different covalent mechanisms^67,68^ while unmodified fluorescein acts as a negative control (Fig. 2C). High fluorescence signals were detected only in microfluidic channels constructed with APTES-coated substrates using amine-reactive probes, confirming the presence of reactive amine groups on the microchannel surfaces (Fig. 2D-E, Fig. S3).

Having confirmed APTES reactivity within sealed microchannels, we sought to photocage surface amines using commercially available, amine-reactive photocleavable PEG (PC PEG) (Fig. 3A). XPS analysis of glass surfaces modified with PC PEG demonstrated a relative increase in carbon and nitrogen content, along with the appearance of a carbonyl peak (Fig. 3B). UV exposure caused a modest decrease in the relative carbon and nitrogen content, but not to levels observed for pristine APTES surfaces (Table S1). We hypothesised that a substantial fraction of PC PEG was either uncleaved or did not dissociate from the surface upon rinsing. To evaluate whether PC PEG photocleavage produced detectable chemical contrasts, we patterned PC PEG with a dibenzocyclooctyne (DBCO) head group within the microfluidic channels. This PC PEG could be labelled with Cy3-azide, and APTES amine groups with NHS-Fluorescein (Fig. 3C, Fig. S4). Cy3-Azide was retained only in microchannels containing PC DBCO; UV exposure led to a ∼70% decrease in fluorescence intensity, indicating the partial removal of PC PEG (Fig. 3D). Moreover, photocleavage yielded a detectable NHS-Fluorescein signal within UV-exposed regions, consistent with amine photodeprotection (Fig. 3D). NHS-Fluorescein was also patterned using PC PEG compounds with different PEG lengths and head groups (biotin and methyl) (Fig. S5). Cy3-azide and NHS-Fluorescein pattern intensity could be controlled through the UV dose (Fig 3E-F). Arbitrary shapes were patterned using a digital micromirror device (DMD) (Fig. 3G).

**Figure 3:**
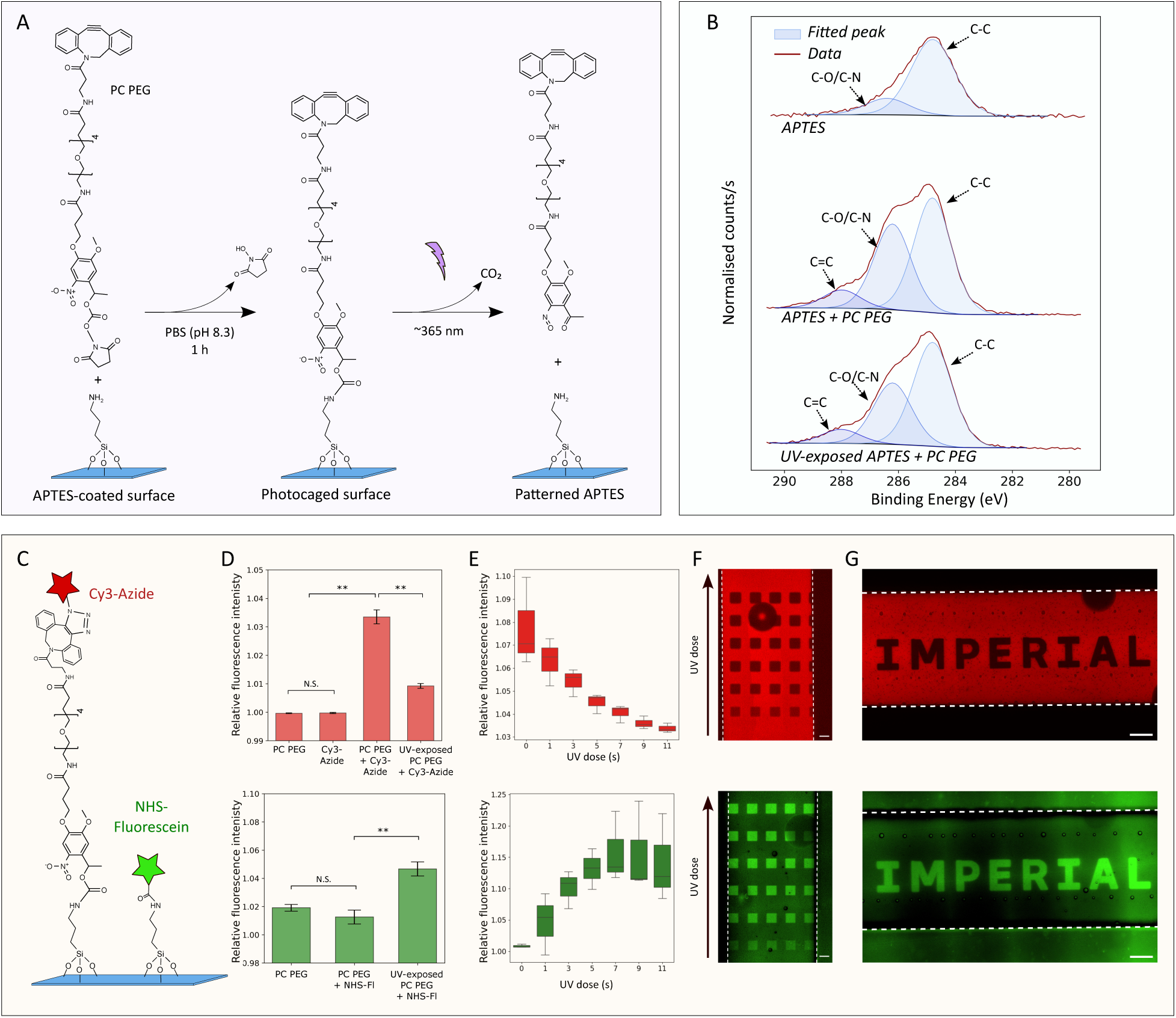
APTES photocaging with PC PEG. **(A)** APTES amine group photocaging and UV-mediated deprotection using PC PEG (with a DBCO head group). **(B)** XPS C1s spectra. PC PEG coating introduced carbonyl peaks; UV exposure slightly reduced carbon content, but did not reproduce the APTES signal. **(C)** PC PEG (with DBCO head group) and APTES labelling with Cy3-Azide and NHS-Fluorescein. **(D)** Pattern quantification: top, Cy3-azide labelling of PC PEG patterns. UV exposure reduced fluorescence by 70% (Kruskal-Wallis and post-hoc Dunn’s test, n=8-16). Bottom: amine groups labelled with NHS-Fluorescein. UV exposure resulted in an increase in fluorescence signal (ANOVA and post-hoc t-test, n=3 for PC PEG; n=6 for other groups). **(E)** Cy3-azide (top) and NHS-Fluorescein intensity in UV-exposed patterns as a function of UV exposure time. Box plots represent the median (line), 25th-75th percentiles (box) and range (whiskers). 1 second corresponds to a UV dose of 62.5 mJ.cm-2 (n=3). **(F)** UV dose effects on Cy3-azide (top) and NHS-Fluorescein (bottom) patterns, with equal UV doses within each row. Increasing UV dose darkens Cy3-azide patterns and brightens NHS-Fluorescein patterns, consistent with increasing levels of amine photodeprotection. Scale bar: 50 µm. **(G)** DMD-printed pattern co-labelled with Cy3-azide (top) and NHS-Fluorescein (bottom). Scale bar: 100 µm.

After characterizing PC PEG and amine patterns, we tested whether deprotected APTES patterns could be functionalized. At neutral pH, APTES surfaces are positively charged.^69^ We first evaluated non-covalent DNA binding to APTES patterns through its negatively charged phosphate backbone (Fig. 4A). Fluorophore-tagged DNA was flown into UV-exposed devices and localised to patterns in both glass-PDMS and PDMS-PDMS device (Fig. 4B-C). We observed no significant difference in pattern relative fluorescence intensity between substrate materials, or between single-stranded and double-stranded DNA (Fig. S6).

**Figure 4:**
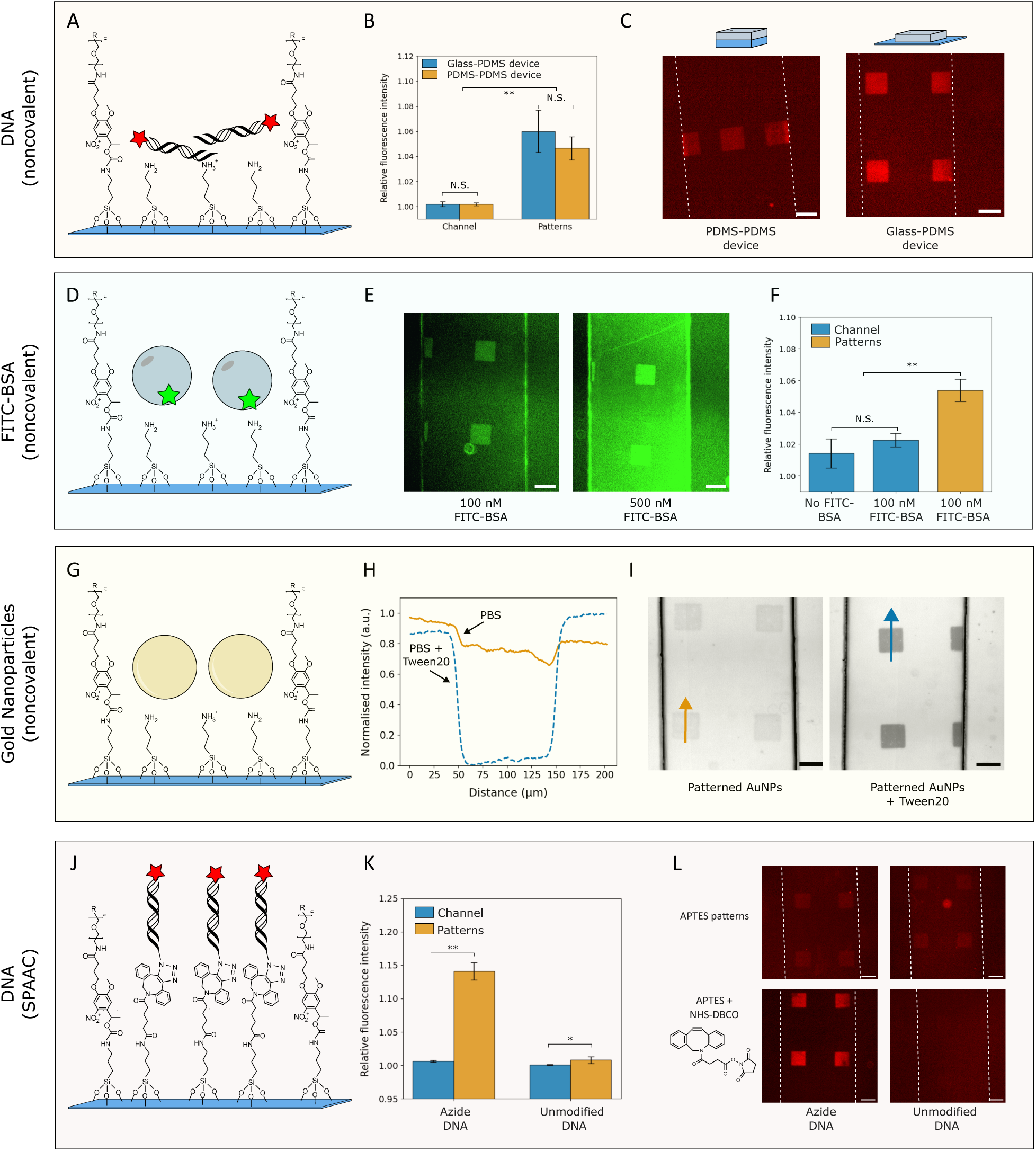
Noncovalent and covalent functionalization of APTES patterns. **(A)** Noncovalent DNA binding to APTES patterns via charge-charge interactions. **(B)** Noncovalent DNA binding on glass and PDMS substrates, with significant differences between pattern and channel fluorescence (n=7-8, Kruskal-Wallis and post-hoc Dunn’s test). **(C)** Representative images of noncovalent DNA patterns formed within PDMS-PDMS and glass-PDMS device using a DMD and mask aligner for UV projection, respectively. Scale bar: 100 µm. **(D)** FITC-BSA noncovalent binding to APTES patterns. **(E)** FITC-BSA concentration dependence: higher concentrations yield significant nonspecific binding outside patterns. **(F)** Channel and pattern fluorescence of 100 nM FITC-BSA patterns. Patterned areas are significantly brighter than unexposed areas (n=4 for no FITC-BSA and n=8 for other groups, Kruskal-Wallis and post-hoc Dunn’s test). **(G)** Noncovalent patterning of gold nanoparticles. **(H)** Line profiles of representative gold nanoparticle patterns; Tween20 increases the signal to noise ratio. **(I)** Representative brightfield images of gold nanoparticle patterns with and without Tween20, arrows indicate regions selected for the line profiles. **(J)** Covalent DNA patterning through SPAAC. After UV exposure, APTES patterns are functionalized with NHS-DBCO, producing ‘clickable’ patterns. Azide-modified DNA is then flown into devices and incubated overnight to form covalent triazole bonds. **(K)** Azide-modified DNA binds APTES + NHS-DBCO patterns, while unmodified DNA does not (n=16 (unmodified DNA) and n=28 (Azide-DNA), Mann-Whitney U test). **(L)** Example images of noncovalent and covalent DNA patterns: NHS-DBCO blocks noncovalent DNA binding while enabling SPAAC-mediated azide-DNA patterns. Scale bars: 100 µm.

We then tested pattern functionalization with proteins. We first evaluated FITC-tagged bovine serum albumin (FITC-BSA) patterning (Fig. 4D). Clear patterns formed at low concentrations (100 nM), whereas higher concentrations led to significant protein adsorption outside of patterns (Fig. 4E-F). ATTO550-conjugated streptavidin, however, was consistently localized outside of patterns, even at low concentrations (Fig. S8). Noncovalent protein patterning on APTES was therefore reliable only at low concentration and dependent on protein-specific interactions.

Lastly, we tested noncovalent pattern functionalization with 20 nm diameter gold nanoparticles, which were expected to have high negative surface charges (Fig. 4G). Gold nanoparticles localized to patterns at densities detectable by optical microscopy, with pattern signals exceeding positive controls (Fig. S7). Interestingly, the addition of the surfactant Tween20 in the gold nanoparticle solution greatly increased the pattern signal-to-noise ratio (Fig. 4H-I).

While functionalizing APTES patterns with noncovalent interactions is simple, it is restricted to negatively charged cargoes and, in theory, would be sensitive to environmental conditions (e.g., charge screening in hypertonic buffers). To increase the versatility of APTES patterns, we generated ‘clickable’ surfaces by patterning surfaces using PC PEG compounds lacking the DBCO head group and flowing NHS-DBCO into devices after UV exposure. DBCO-functionalized patterns could then bind azide-modified cargoes via strain-promoted azide–alkyne cycloaddition (SPAAC) to form covalent triazole linkages (Fig. 4J). After NHS-DBCO conjugation, APTES patterns could no longer be noncovalently bound by unmodified DNA. Conversely, azide-modified DNA produced bright patterns (Fig. 4K-L). SPAAC-bound DNA brushes were denser than noncovalently bound brushes; this effect was also observed in positive controls lacking PC PEG (Fig. S7, S9). While positive tone SPAAC patterns were produced with PC biotin and NHS-DBCO, negative tone patterns could also be produced by clicking azide-modified DNA to UV-patterned PC DBCO (Fig. S10). At NHS-DBCO concentrations above 10 *µ*M, we observed high nonspecific binding outside of intended patterns (Fig. S11). This interaction was mitigated by rinsing microchannels with guanidine hydrochloride (GnHCl), a chaotropic agent (Fig. S12), suggesting a noncovalent mechanism for this unintended adsorption. NHS-Fluorescein was also prone to nonspecific binding at high concentrations (Fig. S13), so we hypothesised that the NHS group may be prone to nonspecific interactions with PC PEG. In addition, we found that patterns were produced only within narrow PC PEG concentration ranges. Low PC PEG concentrations led to high off-pattern binding, presumably because not all APTES amine groups were photocaged. Conversely, at high PC PEG concentrations, patterns became faint or undetectable, possibly indicating the formation of complex PC PEG layers that did not dissociate with the surface after photocleavage. The optimal PC PEG concentration was specific to the PC PEG type and patterned cargo, but did not vary across substrate materials (Fig. S14).

We now consider the behaviour of SPAAC and noncovalently-bound DNA brushes for downstream applications. A motivation to produce covalently bound DNA brushes was to produce heat-resistant bonds, so that anchored single-stranded DNA (ssDNA) strands could be functionalized with complementary ssDNA probes, and recovered by denaturing DNA duplexes using heat, effectively ‘recycling’ patterns (Fig. 5A). As a proof-of-concept, we tested the capture of fluorophore-labelled ssDNA on patterned DNA brushes. Interestingly, while SPAAC-bound patterns readily discriminated between complementary and noncomplementary sequences, noncovalently bound patterns were labelled regardless of the probe sequence (Fig. 5B, Fig. S15). We hypothesized that noncovalent DNA patterns were not dense enough to effectively block direct probe binding to the APTES surface. We were able to denature SPAAC-anchored DNA duplexes, reducing pattern signal below detectable levels. Fresh ssDNA probes subsequently bound the patterns in a sequence-specific manner, suggesting that the initial single-stranded anchors had not dissociated from the surface, as intended (Fig. 5C). As a proof of concept for multi-patterning of different DNA strands, we sequentially immobilized ssDNA anchors with different sequences and incubated a mixed probe ssDNA solution with different fluorophores (Fig. 5D). This approach yielded distinguishable patterns for both SPAAC-functionalized and noncovalently bound DNA, although the latter were consistently fainter (Fig. 5E).

**Figure 5:**
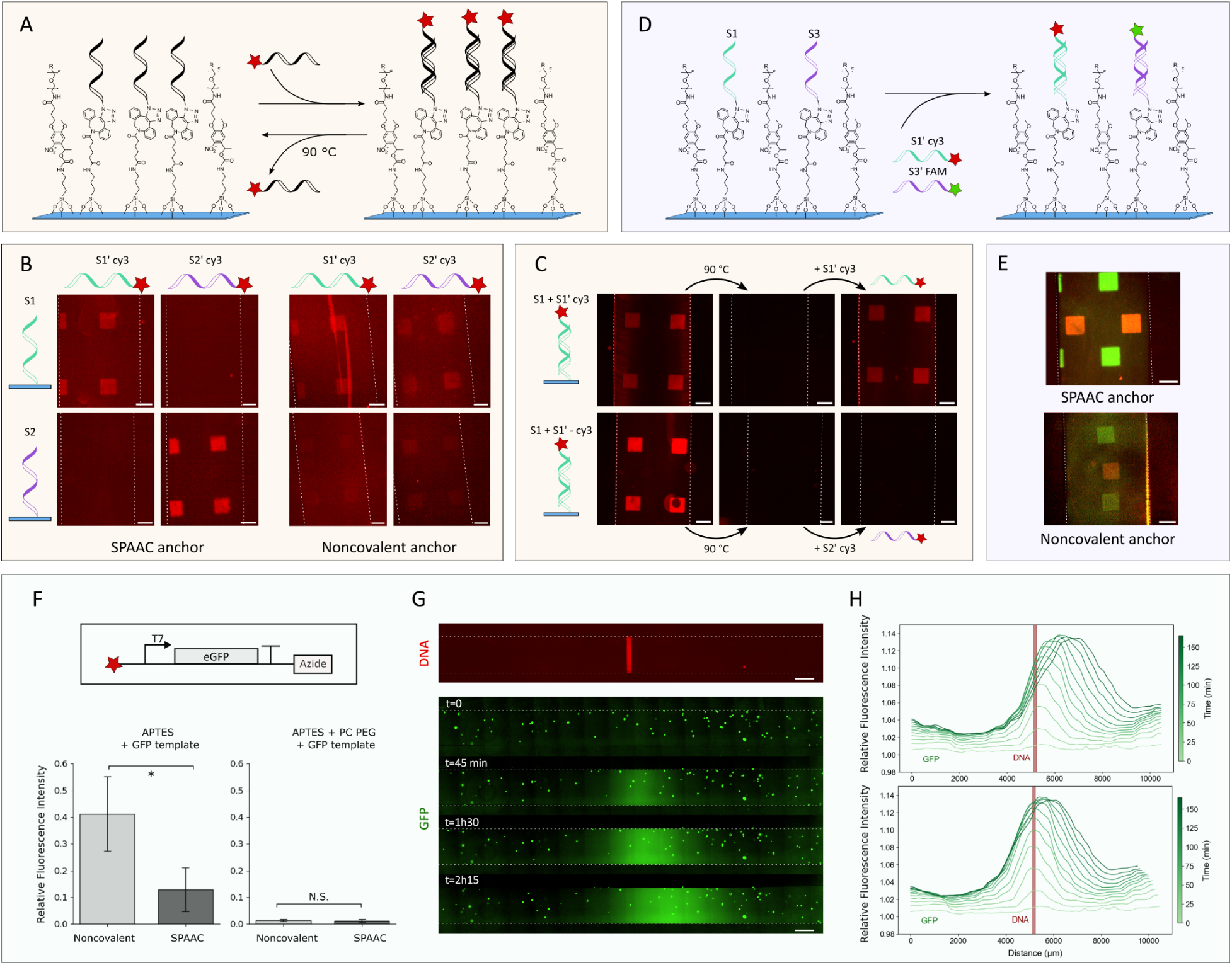
DNA pattern functionalization. **(A)** Single-stranded DNA (ssDNA) capture and heat mediated release from surface-anchored DNA patterns. **(B)** Sequence-selective ssDNA binding on surface-anchored patterns. While SPAAC-bound DNA patterns discriminated between probe sequences, noncovalently bound brushes were faintly labelled, regardless of sequence complementarity. Scale bars: 100 µm. **(C)** Heat mediated release and re-capture of DNA probes. Probe re-capture was sequence specific, suggesting retention of the initial single-stranded brushes. **(D)** Detection of sequentially patterned, single-stranded brushes. **(E)** Proof-of-concept composite images of sequentially patterned DNA with SPAAC and noncovalent anchoring. Scale bars: 100 µm. **(F)** Top: DNA template construct for cellfree GFP expression. Bottom: GFP signal after 1h PURExpress incubation in microfluidic devices containing DNA template bound to APTES or APTES + PC PEG. SPAAC-bound DNA template yielded less GFP (n=6, Mann-Whitney U test). PC PEG coating blocked DNA binding to the surface, leading to no detectable GFP expression (n=8). **(G)** GFP expression from patterned DNA stripe. Top: noncovalently-bound DNA stripe in the centre of the microfluidic channel at t=0. Bottom: GFP expression timelapse, with PURExpress mixed with fluorescent beads (used to track fluid movement) added at t=0. **(H)** Top: GFP signal accross the channel as a function of time. Bottom: GFP signal corrected for fluiddriven drift using fluorescent beads.

Lastly, we evaluated cell-free gene expression from surface-bound DNA. We tested if DNA bound to the surface through covalent or noncovalent binding mechanisms could be transcribed into RNA, and if the patterning method could be used to localise gene expression. We first immobilised the DNA template within microfluidic channels using noncovalent or SPAAC anchoring, without patterning. Microfluidic channels were then rinsed and incubated with a cell-free gene expression mix. While both noncovalent and SPAAC-bound DNA templates produced GFP, we observed higher gene expression levels for noncovalently bound DNA (Fig. 5F, Fig S16). We then repeated the same experiment using microfluidic channels coated with PC PEG prior to DNA incubation. For both SPAAC and noncovalent binding, PC PEG coating reduced GFP expression to levels comparable with controls lacking DNA (Fig. 5F). As a proof-of-concept experiment for localised gene expression, we patterned a 100 µm-wide APTES ‘stripe’ in the centre of a microfluidic channel and immobilised the DNA template via noncovalent binding. While we observed GFP production in the centre of the channel, the peak drifted over time (Fig. S17). We hypothesized that a slow fluid flow occurred within the microfluidic channel, perhaps due to liquid evaporation at the inlets. Consequently, the experiment was repeated using fluorescent polystyrene beads to track fluid flow (Fig. 5G). The GFP peak gradually shifted in the same direction as the subset of polystyrene beads that were observed to be mobile (Fig. 5H). Control microfluidic channels uniformly coated with DNA did not yield a GFP peak (Fig. S18).

## Discussion

We have developed a method to pattern surfaces within microfluidic devices by photolithography using only commercially available reagents and equipment. This method is based on photopatterning APTES amine groups. APTES has been used to coat a range of materials, and mediate surface binding of a range of cargos. For this reason, a patterning method using APTES is versatile: we demonstrated, as a proof of concept, patterning surfaces within microfluidic devices with DNA, proteins, and gold nanoparticles. APTES patterns could be functionalized both covalently and noncovalently. While noncovalent interactions reliably localised DNA strands to patterns, this method was not as effective for proteins due to nonspecific adsorption outside patterns. Gold nanoparticles, however, did non-covalently localize to patterns, although we noticed that Tween20 significantly increased the signal to noise ratio – possibly by reducing charge repulsion between the negatively charged nanoparticles.

DNA and antibodies are often bound to patterns using biotin-streptavidin conjugation.^30,33,40,70,71^ While biotin-streptavidin binding is near-covalent in strength, it can be denatured with heat, and streptavidin can non-specifically adsorb onto surfaces.^72^ Instead of using biotin-streptavidin conjugation, we functionalized APTES with NHS-DBCO, forming versatile ‘clickable’ patterns to covalently bind azide-modified cargoes via SPAAC, a onestep biorthogonal reaction occurring in aqueous conditions.^73,74^ This conjugation method enabled us to denature DNA duplexes (using heat) without disrupting the anchoring mechanism. While SPAAC-bound DNA patterns were denser than noncovalently bound patterns, we found that higher levels of cell-free gene expression occurred from noncovalently bound DNA. We hypothesised that RNA polymerases may be more efficiently released from DNA templates in noncovalent brushes, or that higher DNA density in SPAAC brushes reduced transcription efficiency through crowding effects.^70^

Futher work could improve the robustness of the platform. We found that patterning only worked within narrow PC PEG and cargo concentration ranges. Outside of these ranges, we observed either no detectable binding within patterns, or significant nonspecific adhesion outside patterns. A better understanding of the nature of such off-target interactions would guide mitigation strategies, for example by introducing passivating agents during specific protocol steps.

APTES photopatterning could, in theory, be expanded to different materials. As a proof-of-concept, we demonstrated APTES photopatterning on glass and PDMS surfaces. We expect that any surface that can be coated with APTES can, in theory, be bound to a PDMS top layer and subsequently patterned with DNA, proteins, and gold nanoparticles. In addition, other chemical functionalities could be patterned using this strategy. While we only tested photopatterning using PC PEG compounds functionalized with DBCO and biotin groups, other amine-reactive PC PEG compounds functionalized with maleimide, azide or alkyne head groups are commercially available. Reactive PC PEG head groups could, for example, be used to photorelease surface-bound cargos *in situ*.^62,75^ Another layer of complexity could be achieved by using reconfigurable photoreactive compounds^76^ (instead of PC PEG) to engineer dynamic, light-controlled microfluidic surfaces.

Microfluidic platforms containing localised DNA templates can be harnessed to build artificial, cell-like systems. The microfluidic network can continuously supply cell-free reagents to surface-bound DNA brushes – for instance, to express proteins further regulating gene expression.^77^ Additional complexity could be engineered by continuously expressing cotranscriptionally encoded RNA systems^78,79^ capable of carrying out intricate molecular computation *in situ*.

## Methods

### Microfluidic device fabrication

Patterning experiments were carried out in microfluidic devices containing 7 parallel, independent straight channels. Each channel was 50 *µ*m deep, 500 *µ*m wide and 8 mm long (Fig. S1). On-chip gene expression experiments were carried in single channel devices 80 *µ*m deep, 1 mm wide and 10 mm long.

Silicon master moulds were fabricated by standard SU-8 soft lithography.^80^ Silicon wafers were cleaned with acetone (VWR) and 2-propanol (VWR), exposed to an oxygen plasma (5 min, 45 W). Wafers were then spin-coated (30 s, 3000 rpm) with an SU-8 2050 photoresist (Kayaku) and soft baked to evaporate the solvent (1 min at 65 *^◦^*C, 7 min at 95 *^◦^*C). A polyester film photomask (Microlitho) containing the channel design was placed on top of the wafer and exposed using a mask aligner (UV-KUB 3, Kloe) with a wavelength of 365 nm (i-line) and a UV dose of 3.04 J/cm2. The wafer was then baked (6 min at 95 *^◦^*C) and developed approximately 5 min in (1-Methoxy-2-propyl) acetate (Sigma-Aldrich). The developed wafer was hard baked 15 min at 150 *^◦^*C and silanized with chlorotrimethylsilane (Sigma Aldrich) for 1 h in a vacuum. Channel height was verified using an optic profilometer (Profilm3D, ST Instruments).

PDMS (Sylgard 184) was mixed at a 10:1 ratio of base to curing agent and degassed in a vacuum for 30 min, poured on the silicon master mould and cured at 70 *^◦^*C overnight. The devices were then peeled from the mould, diced and 1 mm inlets punched on the channel extremities.

### APTES coating and device bonding

Unless stated otherwise, all reactions were carried out at room temperature (22-23 *^◦^*C), and reagents purchased from Sigma-Aldrich. Glass coverslips (VWR, borosilicate glass) were cleaned by sonicating 5 min in acetone, 2-propanol, and distilled water (18 MΩ · cm)), and dried at 150 *^◦^*C for 30 min. An APTES stock solution was prepared by mixing v/v 50% methanol (VWR), 2.5% dH2O and 47.5% APTES. This solution was kept 1 h at 4 *^◦^*C before use and stored for up to a week.

To coat glass or PDMS substrates with APTES, a solution of 1% APTES in methanol was prepared from the stock solution. Substrates were exposed to an air plasma (1 min, 50 W) and immediately placed in the 1% APTES solution. After 20 min, the substrates were cleaned by sonicating 3 min in ethanol (VWR), dried with an air gun and baked for 5 min at 100 C to evaporate residual solvents. PDMS top layers were exposed to an air plasma (1 min, 50 W) and immediately bonded to the APTES-coated substrates, using slight pressure to ensure conformal contact between surfaces. The devices were baked for a further 30 min at 100 *^◦^*C.

### X-ray photoelectron spectroscopy

X-ray photoelectron spectroscopy (XPS) measurements were performed on glass coverslips. Samples were prepared according to standard APTES and PC PEG protocols; prior to analysis, they were rinsed with deionized water and dried thoroughly with compressed air. Spectra were acquired on a Thermo Fisher Scientific K-Alpha spectrometer equipped with an Al K*α* source, using a 400 *µ*m X-ray spot and charge neutralization with a flood gun. Data were processed in Avantage (v6.7.0). Peaks were fit with a mixed Gaussian/Lorentzian (pseudo-Voigt) line shape with 30% Lorentzian and 70% Gaussian character, and backgrounds subtracted using the software’s Smart procedure. Binding energies were calibrated to the carbon C1s peak at 284.8 eV.

### APTES detection assays

To test the presence of reactive amine groups within microfluidic channels, 100 nM solutions of Fluorescein, NHS-Fluorescein or FITC were prepared in PBS (pH 7.4) and immediately introduced into microchannels made from APTES-coated substrates. Control microfluidic devices were assembled from surfaces incubated in neat methanol (instead of 1% APTES) for 20 min prior to PDMS bonding. After 2h incubation (without flow), microfluidic channels were rinsed by flushing with PBS and 4 M guanidine hydrochloride (GnHCl) (in dH2O) using 1 ml syringes fitted with connectors.

### APTES photocaging

Reagents and rinsing solutions were manually flushed into devices using 1 ml syringes fitted with connectors; incubation steps occurred in the dark, without fluid flow. PC PEG DBCO (Biosynth), PC PEG biotin and PC PEG methyl (BroadPharm) and were solubilized in 10 mM DMSO stock solutions and stored under inert atmosphere at -20 in the dark. To photocage APTES, PC PEG aliquots were thawed, diluted into PBS (pH 8.3-8.5) and immediately loaded into devices. Unless stated otherwise, PC DBCO, PC biotin and PC methyl concentrations were 100 *µ*M, 2 mM, and 800 *µ*M, respectively. Devices were rinsed with PBS (pH 7.4) after 1h.

### Photopatterning microfluidic devices

UV patterns were projected onto devices using either a mask aligner (UV-KUB 3, Kloe) or a digital micromirror device (DMD) (Smart-Print UV, Microlight3D). To print patterns using a mask aligner, a polyester film photomask (Microlitho) containing arrays of 50 *µ*m wide squares was placed directly on top of devices, using a water drop between the device and the photomask to ensure contact. The devices were then exposed with 72 J/cm^2^. While this method enabled rapid, parallel exposure of multiple devices, it was compatible only with thin substrates (e.g., glass coverslips) as thicker substrates produced out-of-focus patterns. To pattern devices with thicker substrates, the DMD setup was used, using the inbuilt camera to focus on the channel. While slower, the DMD system could be used to print arbitrary patterns and align UV projections to specific microfluidic features. After UV exposure (7.8 J/cm^2^, 385 nm), all devices were rinsed again in PBS (pH 7.4).

### Pattern functionalization

#### NHS-fluorescein and Cy3-Azide

APTES and PC DBCO were tagged with NHS-Fluorescein and Cy3-Azide probes. UV-exposed devices were loaded with 100 nM probes in PBS (pH 7.4) and incubated overnight wrapped in parafilm to reduce evaporation. The devices were then rinsed with 3 times with GnHCl, approximately 5 minutes apart (using 1ml syringes fitted with connectors) and once with PBS.

#### Noncovalent APTES pattern modifications

Patterned devices were loaded with 100 nM DNA in PBS (pH 7.4) and incubated for 1 h. For experiments comparing covalently and non-covalently bound patterns, DNA was incubated overnight. After incubation, devices were rinsed with a 1 M NaCl wash solution. To produce protein patterns, 100 nM FITCBSA was incubated for 1 h and subsequently rinsed in PBS (pH 7.4). Gold nanoparticles (20 nm diameter, 1 OD in suspended in citrate buffer) solutions with and without 1% Tween20 were incubated for 1 h and rinsed with PBS (pH 7.4) prior to imaging.

#### Covalent APTES pattern modification

SPAAC patterns were produced using PC PEG with a biotin head group. UV-exposed devices were loaded with 50 *µ*M NHS-DBCO in PBS (pH 8.3-8.5). After 1 h, the devices were then rinsed 3 times with GnHCl, approximately 5 minutes apart, and once with PBS. Devices were then loaded with 100 nM Azide-modified DNA, wrapped in parafilm, and left to react overnight. Finally, devices were rinsed with a 1 M NaCl wash solution prior to imaging.

#### DNA pattern functionalization

Custom DNA sequences were purchased from Integrated DNA Technologies (Table S2) and solubilized in Tris-EDTA (TE) buffer. Fluorophore and azide-modified strands were ordered with high-performance liquid chromatography purification. DNA duplexes were annealed at 100 *µ*M equimolar concentration in TE + 50 mM NaCl buffer (pH 8.5) by heating to 95 *^◦^*C for 5 min and cooling to 20 *^◦^*C at a rate of 1 *^◦^*C per minute. DNA stock solutions were stored at -20 *^◦^*C.

### DNA pattern hybridization and denaturation

Single-stranded S1 or S2 sequences (Table S2) were patterned using standard covalent and non-covalent patterning protocols. 100 nM solutions of complementary fluorophore-tagged S1’ and S2’ in PBS (pH 7.4) were then loaded into devices. After 2 h, devices were rinsed with 1 M NaCl wash solution. To ‘erase’ patterns, devices were placed on a 90 *^◦^*C hot plate and continuously perfused with PBS (pH 7.4) for 30 min using a syringe pump (AL-1200, WPI) with a flow rate of 2 *µ*l.min-1. Non-erased controls were perfused with PBS at room temperature.

### DNA pattern multiplexing

To produce distinct patterns containing different DNA sequences, single-stranded S1 and S3 sequences were sequentially patterned using standard protocols; a mixed 100 nM Cy3-S1’ and FAM-S3’ solution was then loaded in devices and incubated overnight. Devices were rinsed with 1 M NaCl before imaging.

### GFP expression from surface-bound DNA

The GFP template was inspired by a previously described template^40^ and ordered as a gBlock fragment; its design contained a T7 promoter, GFP template, and a T7 terminator (Table S2). GFP templates were amplified by PCR using azide and fluorophore-modified primers (Q5 high-fidelity PCR kit, NEB), using the manufacturer’s instructions and a 62*^◦^*C melting temperature. PCR products were then purified (Monarch cleanup kit, NEB), and the DNA fragment size verified by agarose gel electrophoresis prior to use. To produce template-coated microfluidic devices, a solution of approximately 5 nM GFP template (in PBS) was loaded into patterned devices. Cell-free gene expression was carried out with PURExpress (NEB) prepared according to the manufacturer’s instructions (omitting the DNA template); mixed with fluorescent polystyrene 1 µm-diameter beads (Sigma-Aldrich) and immediately loaded into devices and imaged at a constant temperature of 37 *^◦^*C. All buffers were autoclaved or sterile filtered prior to use. GFP expression timelapses of patterned devices were processed using BaSiC to reduce shading artifacts and baseline drift. ^81^ GFP expression profiles across the x-axis were plotted using a custom python script. Fluid movement within the microfluidic channel was estimated by tracking bead movement (using nearest-neighbour matching between timepoints); a ‘drift’ value was then calculated for each timepoint as the median distance travelled by beads (excluding static beads). The cumulative drift provided a rough estimate of flow-driven GFP transport, and was used to correct the GFP intensity profiles. To plot GFP intensity profiles, fluorescence peaks caused by beads were localised using Z-score outlier detection and replaced with a local median fluorescence intensity. A Gaussian filter was then applied to further smooth GFP profiles.

### Microscope imaging and data analysis

Microscope images were acquired using a Ti-U inverted fluorescence microscope (Nikon) equipped with a Nikon DS-Qi2 camera, using the NIS-elements software’s inbuilt stitching and shading correction functions. Fluorescence images were captured using TRITC and FITC HC filter sets (AHF Analysentechnik). Image brightness and contrast were manually scaled to visualise patterns. In figures directly comparing images, the same scaling was applied to all images. White dashed lines were manually added to indicate channel edges. To quantify images, areas within UV exposed areas and outside UV exposed areas (referred to as ‘channel’) were manually selected and normalised by dividing to the background signal intensity outside of the microfluidic channel. All histograms represent sample means and error bars standard error of the mean. Data analysis was carried out using custom python scripts; statistical tests were carried out with the scipy.stats module. To compare three or more groups, datasets were first tested for normality and homoscedasticity using Shapiro-Wilk and Levene’s tests, respectively. If datasets passed the normality and equal variance assumptions, differences between conditions were tested with ANOVA, and post-hoc pairwise comparisons carried out with independent t-tests. Conversely, datasets failing to meet the assumptions were tested using the Kruskal-Wallis H test and post-hoc pairwise comparisons were carried out with Dunn’s test. In both cases, Bonferroni correction was applied to correct p-values. Pairwise comparisons were carried out using independent t-tests, or Mann-Whittney U-tests if the assumptions of normality and equal variance were not met. We used a threshold significance level of 0.05 and considered differences to be significant (*) when p≤ 0.05 and very significant (**) for p≤ 0.01.

## Supplementary Information

**Fig. S1:** XPS analysis of open glass surfaces.

**Table S1:** XPS atomic compositions.

**Fig. S2:** 7-lane device used to test patterning methods

**Fig. S3:** Representative images of fluorescein, NHS-Fluorescein and FITC binding assays on PDMS-PDMS devices.

**Fig. S4:** Representative images of NHS-Fluorescein and Cy3-azide patterns used for quantification.

**Fig. S5:** NHS-Fluorescein patterning with alternative PC PEG compounds.

**Fig. S6:** Quantification of noncovalent DNA patterning controls and comparison of single and double-stranded DNA patterns.

**Fig. S7:** Supplementary positive and negative controls.

**Fig. S8:** Streptavidin patterning.

**Fig S9:** Comparison of DNA patterns with and without NHS-DBCO.

**Fig. S10:** Positive and negative DNA patterning with PC DBCO.

**Fig. S11:** Effect of NHS-DBCO concentration on positive SPAAC patterns.

**Fig. S12:** Effect of Gn-HCl rinsing on NHS-Fluorescein and NHS-DBCO.

**Fig. S13:** Effect of cargo concentrations.

**Fig. S14:** PC PEG concentration effects.

**Fig. S15:** ssDNA capture on pattern quantification.

**Fig S16:** GFP expression as a function of time.

**Fig. S17:** GFP expression from patterned DNA (without beads).

**Fig. S18:** Example of GFP expression from DNA immobilized in the entire channel.

**Table S2:** DNA sequences

## Acknowledgements

This work was funded by the Engineering and Physical Sciences Research Council (EPSRC) Centre for Doctoral Training in BioDesign Engineering (grand number EP/S022856/1) and the Royal Society (grant number URF11020).

## Data availability

Raw data and processing scripts are available on Zenodo: 10.5281/zenodo.19398836

## Notes

### Competing Interest Statement

The authors have declared no competing interest.

### Summary of Updates

Revised figure 4, additional supplementary materials.

## References

1. Manz, A.; Graber, N.; Widmer, H. M. Miniaturized total chemical analysis systems: A novel concept for chemical sensing. Sensors and Actuators B: Chemical 1990, 1, 244– 248.

2. Prangemeier, T.; Lehr, F. X.; Schoeman, R. M.; Koeppl, H. Microfluidic platforms for the dynamic characterisation of synthetic circuitry. Current Opinion in Biotechnology 2020, 63, 167–176.

3. Allard, P.; Papazotos, F.; Potvin-Trottier, L. Microfluidics for long-term single-cell time-lapse microscopy: Advances and applications. Frontiers in Bioengineering and Biotechnology 2022, 10.

4. Squires, T. M.; Quake, S. R. Microfluidics: Fluid physics at the nanoliter scale. Reviews of Modern Physics 2005, 77, 977–1026.

5. Whitesides, G. M. The origins and the future of microfluidics. Nature 2006, 442, 368– 373.

6. Shakeri, A.; Jarad, N. A.; Leung, A.; Soleymani, L.; Didar, T. F. Biofunctionalization of Glass- and Paper-Based Microfluidic Devices: A Review. Advanced Materials Interfaces 2019, 6.

7. Ashok, D.; Singh, J.; Howard, H. R.; Cottam, S.; Waterhouse, A.; Bilek, M. M. M. Interfacial engineering for biomolecule immobilisation in microfluidic devices. Biomaterials 2025, 316, 123014.

8. Trantidou, T.; Elani, Y.; Parsons, E.; Ces, O. Hydrophilic surface modification of PDMS for droplet microfluidics using a simple, quick, and robust method via PVA deposition. Microsystems & Nanoengineering 2017, 3, 16091.

9. Kazoe, Y.; Ugajin, T.; Ohta, R.; Mawatari, K.; Kitamori, T. Parallel multiphase nanofluidics utilizing nanochannels with partial hydrophobic surface modification and application to femtoliter solvent extraction. Lab on a Chip 2019, 19, 3844–3852.

10. Garcia-Cordero, J. L.; Maerkl, S. J. Multiplexed surface micropatterning of proteins with a pressure-modulated microfluidic button-membrane. Chemical Communications 2013, 49, 1264–1266.

11. Larsen, E. K. U.; Mikkelsen, M. B. L.; Larsen, N. B. Protein and cell patterning in closed polymer channels by photoimmobilizing proteins on photografted poly(ethylene glycol) diacrylate. Biomicrofluidics 2014, 8, 064127.

12. Takayama, S.; McDonald, J. C.; Ostuni, E.; Liang, M. N.; Kenis, P. J. A.; Ismagilov, R. F.; Whitesides, G. M. Patterning cells and their environments using multiple laminar fluid flows in capillary networks. Proceedings of the National Academy of Sciences 1999, 96, 5545–5548.

13. Siddique, A.; Meckel, T.; Stark, R. W.; Narayan, S. Improved cell adhesion under shear stress in PDMS microfluidic devices. Colloids and Surfaces B: Biointerfaces 2017, 150, 456–464.

14. Ahn, S. I.; Sei, Y. J.; Park, H.-J.; Kim, J.; Ryu, Y.; Choi, J. J.; Sung, H.-J.; MacDonald, T. J.; Levey, A. I.; Kim, Y. Microengineered human blood–brain barrier platform for understanding nanoparticle transport mechanisms. Nature Communications 2020, 11, 175.

15. Khodayari Bavil, A.; Kim, J. A capillary flow-driven microfluidic system for microparticle-labeled immunoassays. The Analyst 2018, 143, 3335–3342.

16. Launiere, C.; Gaskill, M.; Czaplewski, G.; Myung, J. H.; Hong, S.; Eddington, D. T. Channel Surface Patterning of Alternating Biomimetic Protein Combinations for Enhanced Microfluidic Tumor Cell Isolation. Analytical Chemistry 2012, 84, 4022–4028.

17. Pardatscher, G.; Schwarz-Schilling, M.; Daube, S. S.; Bar-Ziv, R. H.; Simmel, F. C. Gene Expression on DNA Biochips Patterned with Strand-Displacement Lithography. Angewandte Chemie International Edition 2018, 57, 4783–4786.

18. Sathish, S.; Ricoult, S. G.; Toda-Peters, K.; Shen, A. Q. Microcontact printing with aminosilanes: creating biomolecule micro- and nanoarrays for multiplexed microfluidic bioassays. The Analyst 2017, 142, 1772–1781.

19. Zheng, C.; Wang, J.; Pang, Y.; Wang, J.; Li, W.; Ge, Z.; Huang, Y. High-throughput immunoassay through in-channel microfluidic patterning. Lab on a Chip 2012, 12, 2487– 2490.

20. Tong, Z.; Ivask, A.; Guo, K.; McCormick, S.; Lombi, E.; Priest, C.; Voelcker, N. H. Crossed flow microfluidics for high throughput screening of bioactive chemical–cell interactions. Lab on a Chip 2017, 17, 501–510.

21. Grant, J.; Modica, J. A.; Roll, J.; Perkovich, P.; Mrksich, M. An Immobilized Enzyme Reactor for Spatiotemporal Control over Reaction Products. Small 2018, 14, 1800923.

22. Obst, F.; Simon, D.; Mehner, P. J.; Neubauer, J. W.; Beck, A.; Stroyuk, O.; Richter, A.; Voit, B.; Appelhans, D. One-step photostructuring of multiple hydrogel arrays for compartmentalized enzyme reactions in microfluidic devices. Reaction Chemistry & Engineering 2019, 4, 2141–2155.

23. Vong, T.; Schoffelen, S.; Dongen, S. F. M. v.; Beek, T. A. v.; Zuilhof, H.; Hest, J. C. M. v. A DNA-based strategy for dynamic positional enzyme immobilization inside fused silica microchannels. Chemical Science 2011, 2, 1278–1285.

24. Kosaka, T.; Yamaguchi, S.; Izuta, S.; Yamahira, S.; Shibasaki, Y.; Tateno, H.; Okamoto, A. Bioorthogonal Photoreactive Surfaces for Single-Cell Analysis of Intercellular Communications. Journal of the American Chemical Society 2022, 144, 17980–17988.

25. Renberg, B.; Sato, K.; Mawatari, K.; Idota, N.; Tsukahara, T.; Kitamori, T. Serial DNA immobilization in micro- and extended nanospace channels. Lab on a Chip 2009, 9, 1517–1523.

26. Kim, D. W.; Rubanov, M.; Grinthal, A.; Moerman, P.; Schulman, R. Building RNA concentration fields. Matter 2025, 8, 102208.

27. Tayar, A. M.; Karzbrun, E.; Noireaux, V.; Bar-Ziv, R. H. Propagating gene expression fronts in a one-dimensional coupled system of artificial cells. Nature Physics 2015, 11, 1037–1041.

28. Collins, K.; Stanley, C. E.; Ouldridge, T. E. Biochemical surface patterning in microfluidic devices. Current Opinion in Biotechnology 2025, 96, 103390.

29. Didar, T. F.; Foudeh, A. M.; Tabrizian, M. Patterning Multiplex Protein Microarrays in a Single Microfluidic Channel. Analytical Chemistry 2012, 84, 1012–1018.

30. Gerber, D.; Maerkl, S. J.; Quake, S. R. An in vitro microfluidic approach to generating protein-interaction networks. Nature methods 2009, 6, 71–74.

31. Buxboim, A.; Bar-Dagan, M.; Frydman, V.; Zbaida, D.; Morpurgo, M.; Bar-Ziv, R. A Single-Step Photolithographic Interface for Cell-Free Gene Expression and Active Biochips. Small 2007, 3, 500–510.

32. Feyssa, B.; Liedert, C.; Kivimaki, L.; Johansson, L.-S.; Jantunen, H.; Hakalahti, L. Patterned Immobilization of Antibodies within Roll-to-Roll Hot Embossed Polymeric Microfluidic Channels. PLOS ONE 2013, 8, e68918.

33. Bavli, D.; Ezra, E.; Kitsberg, D.; Vosk-Artzi, M.; Murthy, S. K.; Nahmias, Y. One step antibody-mediated isolation and patterning of multiple cell types in microfluidic devices. Biomicrofluidics 2016, 10, 024112.

34. Loessberg-Zahl, J.; Beumer, J.; van den Berg, A.; Eijkel, J. C. T.; van der Meer, A. D. Patterning Biological Gels for 3D Cell Culture inside Microfluidic Devices by Local Surface Modification through Laminar Flow Patterning. Micromachines 2020, 11, 1112, Number: 12.

35. Delamarche, E.; Pereiro, I.; Kashyap, A.; Kaigala, G. V. Biopatterning: The Art of Patterning Biomolecules on Surfaces. Langmuir 2021, 37, 9637–9651.

36. Rubanov, M.; Cole, J.; Lee, H.-J.; Cordova, L. G. S.; Chen, Z.; Gonzalez, E.; Schulman, R. Multi-domain automated patterning of DNA-functionalized hydrogels. PLOS ONE 2024, 19, e0295923.

37. Böcherer, D.; Li, Y.; Rein, C.; Franco Corredor, S.; Hou, P.; Helmer, D. High-Resolution 3D Printing of Dual-Curing Thiol-Ene/Epoxy System for Fabrication of Microfluidic Devices for Bioassays. Advanced Functional Materials 2024, 34, 2401516.

38. Hynes, M. J.; Maurer, J. A. Lighting the path: photopatternable substrates for biological applications. Mol. BioSyst. 2013, 9, 559–564.

39. Kaneko, S.; Nakayama, H.; Yoshino, Y.; Fushimi, D.; Yamaguchi, K.; Horiike, Y.; Nakanishi, J. Photocontrol of cell adhesion on amino-bearing surfaces by reversible conjugation of poly(ethylene glycol) via a photocleavable linker. Physical Chemistry Chemical Physics 2011, 13, 4051–4059.

40. Pardatscher, G.; Schwarz-Schilling, M.; Sagredo, S.; Simmel, F. C. Functional Surface-immobilization of Genes Using Multistep Strand Displacement Lithography. Journal of Visualized Experiments 2018,

41. Alazzam, A.; Alamoodi, N. Microfluidic Devices with Patterned Wettability Using Graphene Oxide for Continuous Liquid–Liquid Two-Phase Separation. ACS Applied Nano Materials 2020, 3, 3471–3477.

42. Holden, M. A.; Jung, S.-Y.; Cremer, P. S. Patterning Enzymes Inside Microfluidic Channels via Photoattachment Chemistry. Analytical Chemistry 2004, 76, 1838–1843.

43. Castellana, E. T.; Kataoka, S.; Albertorio, F.; Cremer, P. S. Direct Writing of Metal Nanoparticle Films Inside Sealed Microfluidic Channels. Analytical Chemistry 2006, 78, 107–112.

44. Morikawa, K.; Kazumi, H.; Tsuyama, Y.; Ohta, R.; Kitamori, T. Surface Patterning of Closed Nanochannel Using VUV Light and Surface Evaluation by Streaming Current. Micromachines 2021, 12, 1367, Number: 11.

45. Vlachopoulou, M.-E.; Tserepi, A.; Pavli, P.; Argitis, P.; Sanopoulou, M.; Misiakos, K. A low temperature surface modification assisted method for bonding plastic substrates. Journal of Micromechanics and Microengineering 2008, 19, 015007.

46. Kim, K.; Park, S. W.; Yang, S. S. The optimization of PDMS-PMMA bonding process using silane primer. BioChip Journal 2010, 4, 148–154.

47. Lee, M.; Lopez-Martinez, M. J.; Baraket, A.; Zine, N.; Esteve, J.; Plaza, J. A.; Jaffrezic-Renault, N.; Errachid, A. Polymer micromixers bonded to thermoplastic films combining soft-lithography with plasma and aptes treatment processes. Journal of Polymer Science Part A: Polymer Chemistry 2013, 51, 59–70.

48. Sunkara, V.; Park, D.-K.; Hwang, H.; Chantiwas, R.; Soper, S. A.; Cho, Y.-K. Simple room temperature bonding of thermoplastics and poly(dimethylsiloxane). Lab Chip 2011, 11, 962–965.

49. Aran, K.; Sasso, L. A.; Kamdar, N.; Zahn, J. D. Irreversible, direct bonding of nanoporous polymer membranes to PDMS or glass microdevices. Lab on a Chip 2010, 10, 548–552.

50. Sunkara, V.; Park, D.-K.; Cho, Y.-K. Versatile method for bonding hard and soft materials. RSC Advances 2012, 2, 9066–9066.

51. Asadishad, B.; Ghoshal, S.; Tufenkji, N. Method for the direct observation and quantification of survival of bacteria attached to negatively or positively charged surfaces in an aqueous medium. Environmental Science and Technology 2011, 45, 8345–8351.

52. Wang, L.; Feng, X.; Hou, S.; Chan, Q.; Qin, M. Microcontact printing of multiproteins on the modified mica substrate and study of immunoassays. Surface and Interface Analysis 2006, 38, 44–50, eprint: https://onlinelibrary.wiley.com/doi/pdf/10.1002/sia.2178.

53. Sivagnanam, V.; Song, B.; Vandevyver, C.; Bünzli, J.-C. G.; Gijs, M. A. M. Selective Breast Cancer Cell Capture, Culture, and Immunocytochemical Analysis Using Self-Assembled Magnetic Bead Patterns in a Microfluidic Chip. Langmuir 2010, 26, 6091– 6096.

54. Jeon, E.; Koo, B.; Kim, S.; Kim, J.; Yu, Y.; Jang, H.; Lee, M.; Kim, S.-H.; Kang, T.; Kim, S. K.; Kwak, R.; Shin, Y.; Lee, J. Biporous silica nanostructure-induced nanovortex in microfluidics for nucleic acid enrichment, isolation, and PCR-free detection. Nature Communications 2024, 15, 1366.

55. Rahman, M.; Norton, M. L. Two-Dimensional Materials as Substrates for the Development of Origami-Based Bionanosensors. IEEE Transactions on Nanotechnology 2010, 9, 539–542, Conference Name: IEEE Transactions on Nanotechnology.

56. Jin, L.; Horgan, A.; Levicky, R. Preparation of end-tethered DNA monolayers on siliceous surfaces using heterobifunctional cross-linkers. Langmuir 2003, 19, 6968–6975.

57. Pivetal, J.; Pereira, F. M.; Barbosa, A. I.; Castanheira, A. P.; Reis, N. M.; Edwards, A. D. Covalent immobilisation of antibodies in Teflon-FEP microfluidic devices for the sensitive quantification of clinically relevant protein biomarkers. Analyst 2017, 142, 959–968.

58. Wang, A.-J.; Feng, J.-J.; Fan, J. Covalent modified hydrophilic polymer brushes onto poly(dimethylsiloxane) microchannel surface for electrophoresis separation of amino acids. Journal of Chromatography A 2008, 1192, 173–179.

59. Gunda, N. S. K.; Singh, M.; Norman, L.; Kaur, K.; Mitra, S. K. Optimization and characterization of biomolecule immobilization on silicon substrates using (3-aminopropyl)triethoxysilane (APTES) and glutaraldehyde linker. Applied Surface Science 2014, 305, 522–530.

60. Syga,; Spakman, D.; Punter, C. M.; Poolman, B. Method for immobilization of living and synthetic cells for high-resolution imaging and single-particle tracking. Scientific Reports 2018, 8, 13789–13789.

61. Laborie, E.; Bayle, F.; Bouville, D.; Smadja, C.; Dufour-Gergam, E.; Ammar, M. Surface Biochemical Modification of Poly(dimethylsiloxane) for Specific Immune Cytokine Response. ACS Applied Bio Materials 2021, 4, 1307–1318.

62. Yamaguchi, S.; Takasaki, Y.; Yamahira, S.; Nagamune, T. Photo-Cleavable peptide-poly(Ethylene Glycol) conjugate surfaces for light-guided control of cell adhesion. Micromachines 2020, 11.

63. Willems, S. B. J.; Zegers, J.; Bunschoten, A.; Wagterveld, R. M.; Leeuwen, F. W. B. v.; Velders, A. H.; Saggiomo, V. COvalent monolayer patterns in Microfluidics by PLasma etching Open Technology – COMPLOT. Analyst 2020, 145, 1629–1635.

64. Kim, J.; Jensen, E. C.; Megens, M.; Boser, B.; Mathies, R. A. Integrated microfluidic bioprocessor for solid phase capture immunoassays. Lab on a Chip 2011, 11, 3106–3112.

65. Jang, K.; Xu, Y.; Tanaka, Y.; Sato, K.; Mawatari, K.; Konno, T.; Ishihara, K.; Kitamori, T. Single-cell attachment and culture method using a photochemical reaction in a closed microfluidic system. Biomicrofluidics 2010, 4, 032208.

66. Bertrand, O.; Gohy, J.-F. Photo-responsive polymers: synthesis and applications. Polymer Chemistry 2017, 8, 52–73.

67. Kirley, T. L.; Norman, A. B. Characterization and optimization of fluorescein isothiocyanate labeling of humanized h2E2 anti-cocaine mAb. Biochemistry and Biophysics Reports 2023, 35, 101520.

68. Nanda, J. S.; Lorsch, J. R. Labeling a protein with fluorophores using NHS ester derivitization. Methods in Enzymology 2014, 536, 87–94.

69. Sun, Y.; Yanagisawa, M.; Kunimoto, M.; Nakamura, M.; Homma, T. Depth profiling of APTES self-assembled monolayers using surface-enhanced confocal Raman microspectroscopy. Spectrochimica Acta Part A: Molecular and Biomolecular Spectroscopy 2017, 184, 1–6.

70. Buxboim, A.; Daube, S. S.; Bar-Ziv, R. Ultradense Synthetic Gene Brushes on a Chip. Nano Letters 2009, 9, 909–913.

71. Eyer, K.; Kuhn, P.; Stratz, S.; Dittrich, P. S. A Microfluidic Chip for the Versatile Chemical Analysis of Single Cells. Journal of Visualized Experiments (JoVE) 2013, e50618.

72. Ylikotila, J.; Välimaa, L.; Takalo, H.; Pettersson, K. Improved surface stability and biotin binding properties of streptavidin coating on polystyrene. Colloids and Surfaces B: Biointerfaces 2009, 70, 271–277.

73. Agard, N. J.; Prescher, J. A.; Bertozzi, C. R. A Strain-Promoted [3 + 2] AzideAlkyne Cycloaddition for Covalent Modification of Biomolecules in Living Systems. Journal of the American Chemical Society 2004, 126, 15046–15047.

74. Jewett, J. C.; Bertozzi, C. R. Cu-free click cycloaddition reactions in chemical biology. Chemical Society Reviews 2010, 39, 1272–1279.

75. Usman, A.; Asghar Eftekhar, A.; Adibi, A. Affinity-Based Capturing, Release, and Glycoprofiling of PSA Cancer Biomarker Using Miniaturized Micropillar-Based Platform. IEEE Sensors Journal 2024, 24, 17395–17402, Conference Name: IEEE Sensors Journal.

76. Liu, J.; Butt, H.-J.; Wu, S. Reconfigurable Surfaces Based on Photocontrolled Dynamic Bonds. Advanced Functional Materials 2020, 30, 1907605.

77. Karzbrun, E.; Tayar, A. M.; Noireaux, V.; Bar-Ziv, R. H. Programmable on-chip DNA compartments as artificial cells. Science 2014, 345, 829–832.

78. Samuel W Schaffter; Strychalski, E. A. Cotranscriptionally encoded RNA strand displacement circuits; 2022; pp 4354–4354, Volume: 8.

79. Bae, W.; Stan, G.-B. V.; Ouldridge, T. E. In situ Generation of RNA Complexes for Synthetic Molecular Strand-Displacement Circuits in Autonomous Systems. Nano Letters 2021, 21, 265–271.

80. Qin, D.; Xia, Y.; Whitesides, G. M. Soft lithography for micro- and nanoscale patterning. Nature Protocols 2010, 5, 491–502.

81. Peng, T.; Thorn, K.; Schroeder, T.; Wang, L.; Theis, F. J.; Marr, C.; Navab, N. A BaSiC tool for background and shading correction of optical microscopy images. Nature Communications 2017, 8, 14836.

