## Supplementary Materials for "A versatile method to pattern surfaces within microfluidic devices"

### Table of Contents

**Fig. S1:** XPS analysis of open glass surfaces.

**Table S1:** XPS atomic compositions.

**Fig. S2:** 7-lane device used to test patterning methods.

**Fig. S3:** Representative images of fluorescein, NHS-Fluorescein and FITC binding assays on PDMS-PDMS devices.

**Fig. S4:** Representative images of NHS-Fluorescein and Cy3-azide patterns used for quantification.

**Fig. S5:** NHS-Fluorescein patterning with alternative PC PEG compounds.

**Fig. S6:** Quantification of noncovalent DNA patterning controls and comparison of single and double-stranded DNA patterns.

**Fig. S7:** Supplementary positive and negative controls.

**Fig. S8:** Streptavidin patterning.

**Fig. S9:** Comparison of DNA patterns with and without NHS-DBCO.

**Fig. S10:** Positive and negative DNA patterning with PC DBCO.

**Fig. S11:** Effect of NHS-DBCO concentration on positive SPAAC patterns.

**Fig. S12:** Effect of Gn-HCl rinsing on NHS-Fluorescein and NHS-DBCO.

**Fig. S13:** Effect of cargo concentrations.

**Fig. S14:** PC PEG concentration effects.

**Fig. S15:** ssDNA capture on pattern quantification.

**Fig. S16:** GFP expression as a function of time.

**Fig. S17:** GFP expression from patterned DNA (without beads).

**Fig. S18:** Example of GFP expression from DNA immobilized in the entire channel.

**Table S2:** DNA sequences.

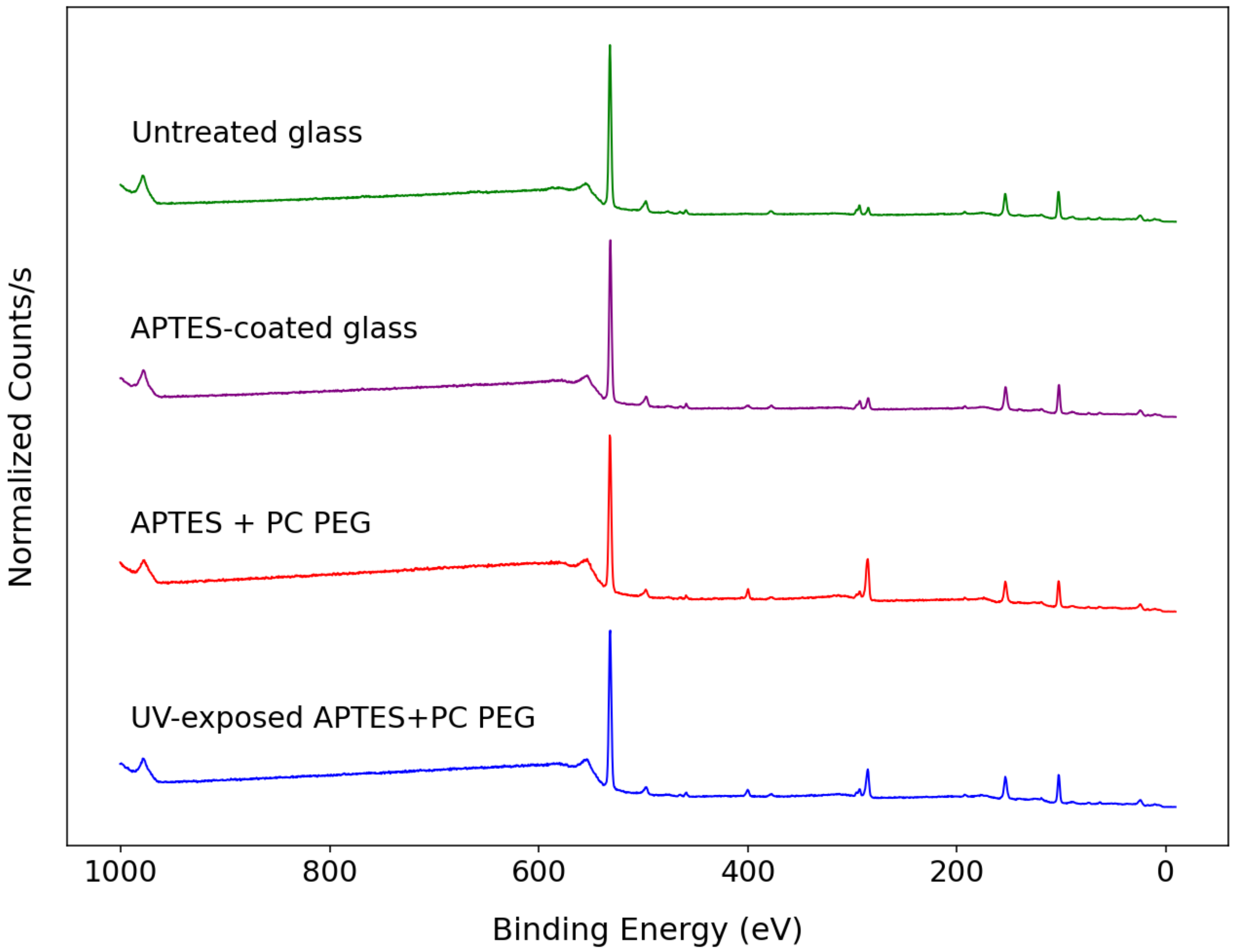


**Figure S1**: XPS analysis of modified glass surfaces. APTES coating led to an N1s peak around 400 eV, PC PEG coating led to an increased C1s peak around 300 eV. Other notable peaks include O1s ($\approx$ 550 eV) and Si2p ($\approx$ 100 eV).

**Table S1:** XPS atomic composition (at. %) calculated from background-subtracted peak areas. Atomic compositions reflect the elemental composition at the surface of the sample. APTES and PC PEG addition led to an increase in carbon and nitrogen content, which was partially reduced by UV exposure.

| Sample | C (at.%) | N (at.%) | Si (at.%) | O (at.%) |
|---|---|---|---|---|
| Untreated glass | 2.98 | 0.32 | 38.78 | 57.92 |
| APTES-coated glass | 7.8 | 1.44 | 23.13 | 67.63 |
| APTES + PC PEG | 29.27 | 4.07 | 16.37 | 50.29 |
| UV-exposed APTES + PC PEG | 20.23 | 3.03 | 19.31 | 57.42 |

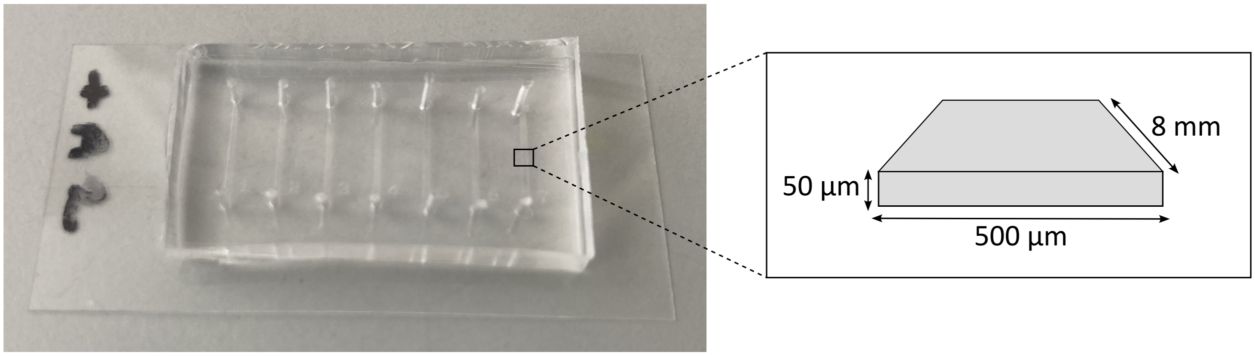


**Figure S2:** 7-lane device used to test patterning methods. Left: glass-PDMS device. The glass coverslip was coated with APTES, and bound to the plasma-treated PDMS top layer to form microchannels. Right: channel dimensions.

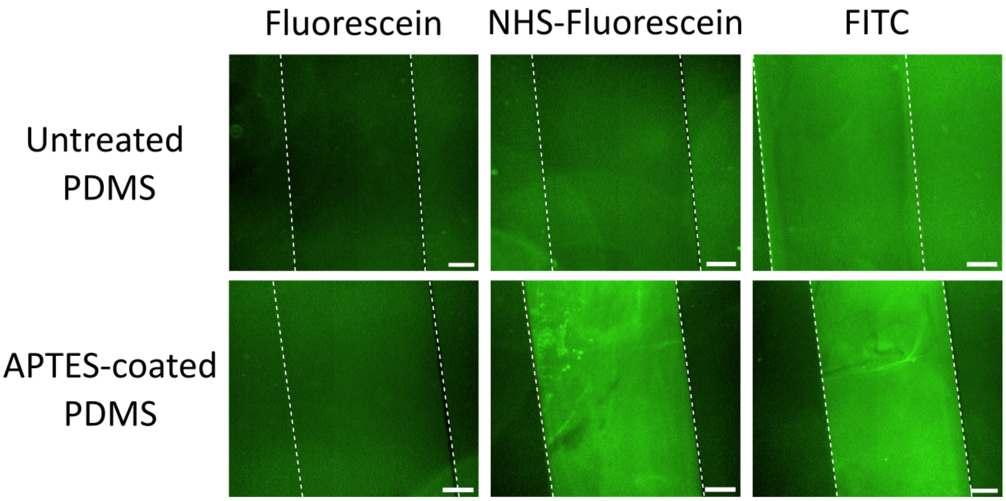


**Figure S3:** Representative images of fluorescein, NHS-Fluorescein and FITC binding assays on PDMS-PDMS devices. APTES-coated PDMS-PDMS devices were manufactured by binding an APTES-coated substrate layer to a plasma-exposed top layer. An increase in channel fluorescence was observed only in samples containing APTES-coated PDMS substrate layers and using amine reactive probes, indicating that APTES coating produced accessible surface amine groups. Dashed white lines represent channel edges; images were scaled to the same brightness and contrast settings for direct comparison. Scale bar: 100 µm.

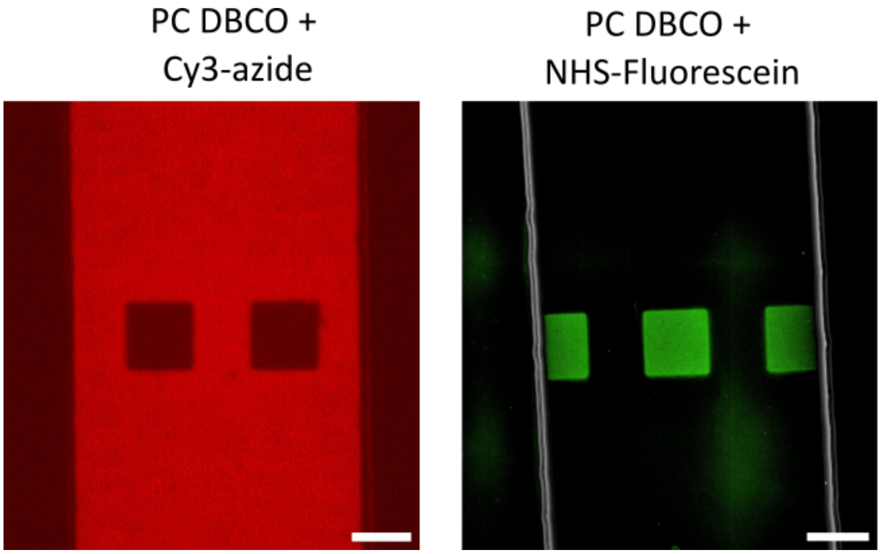


**Figure S4:** Representative images of Cy3-azide (right) and NHS-Fluorescein (left) patterns used for quantification. 100 µm-wide patterns were projected using a DMD, images were acquired using a 20x microscope objective. Scale bar: 100 µm.

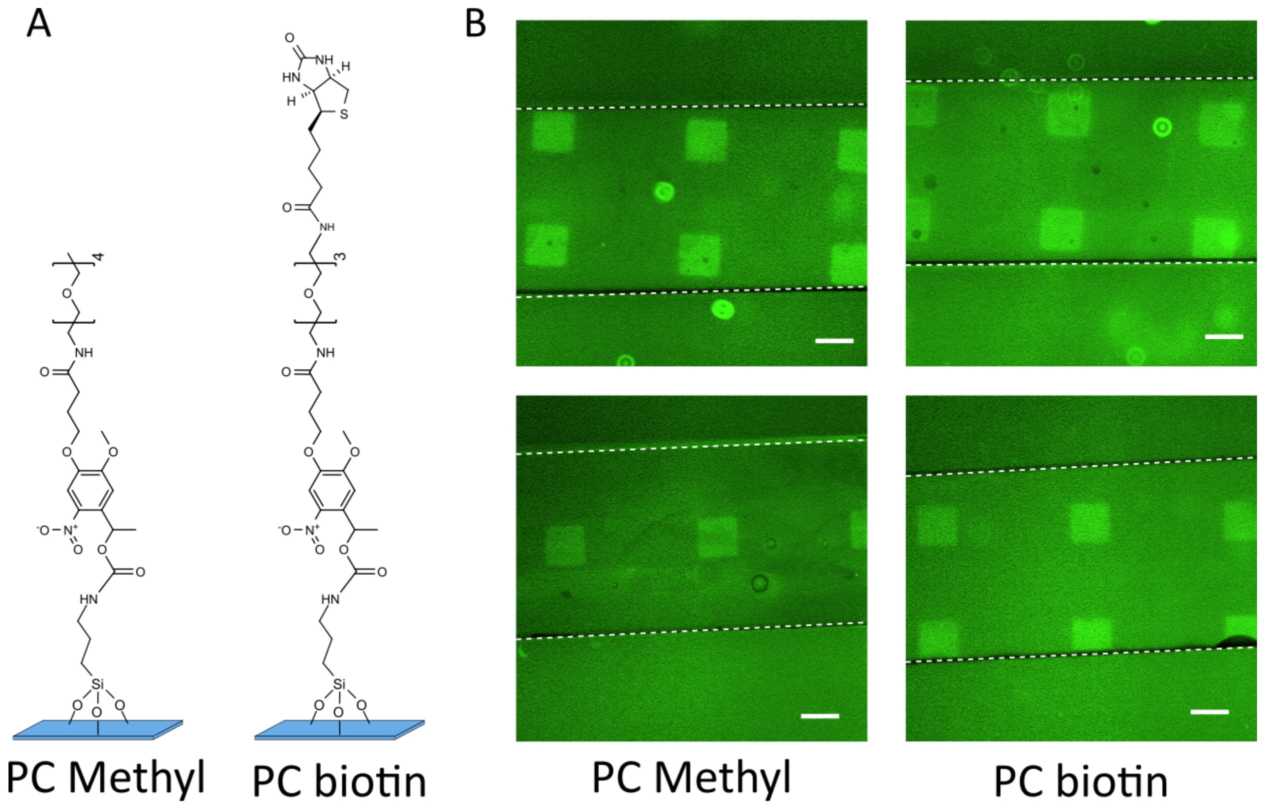


**Figure S5:** NHS-Fluorescein patterning with alternative PC PEG compounds. **(A)** Chemical structure PC methyl and PC biotin bound to APTES surfaces. **(B)** Example microscope images of NHS-Fluorescein patterned in glass-PDMS devices using PC methyl and PC biotin. UV patterns were exposed onto PC PEG-coated microfludic channels using a mask aligner. Scale bar: 100 µm.

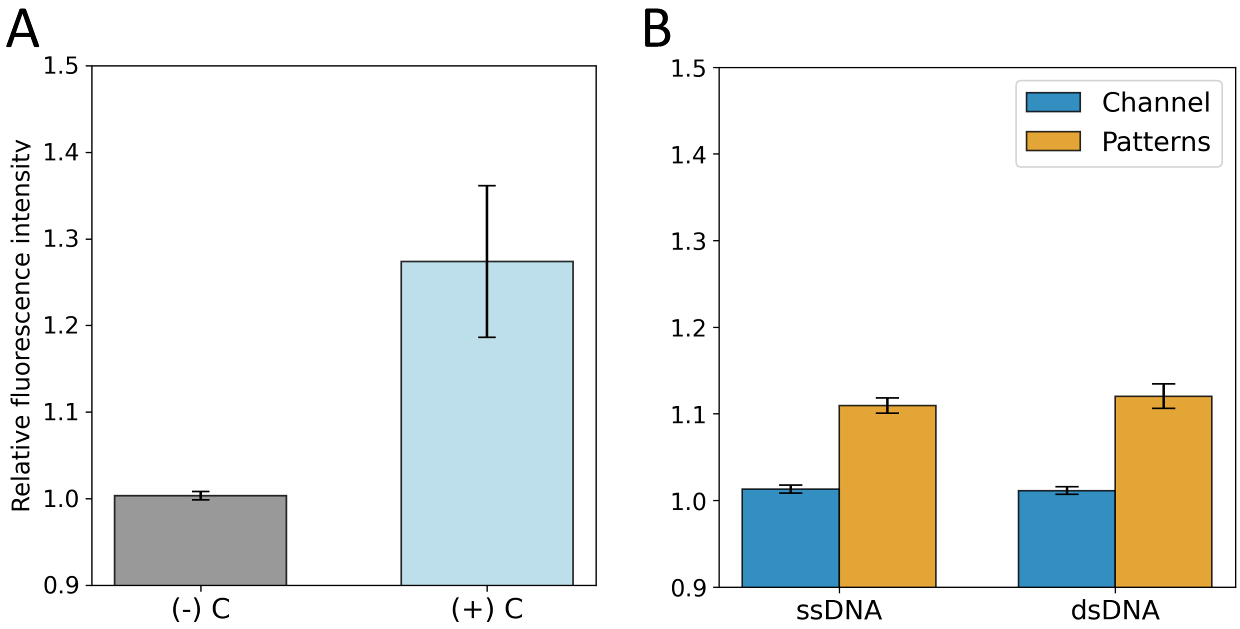


**Figure S6:** Supplementary quantification of noncovalent DNA patterns. **(A)** Comparison of negative and positive controls. Negative controls were channels coated with PC PEG without adding DNA; positive controls contained no PC PEG, so DNA could bind the entire APTES-modified surface. The fluorescence intensity within channels was normalised by dividing with the background fluorescence intensity outside channels (n=5). **(B)** Comparison of patterns produced using single stranded and double stranded DNA probes. Devices were incubated with PC PEG, UV exposed using a mask aligner (printing 50 µm wide square patterns), rinsed, and incubated with either single-stranded DNA (n=7) or double-stranded DNA (n=9). Pattern and channel fluorescence was normalised to background fluorescence outside microchannels; 'channel' denotes areas outside patterns, which were not UV exposed.

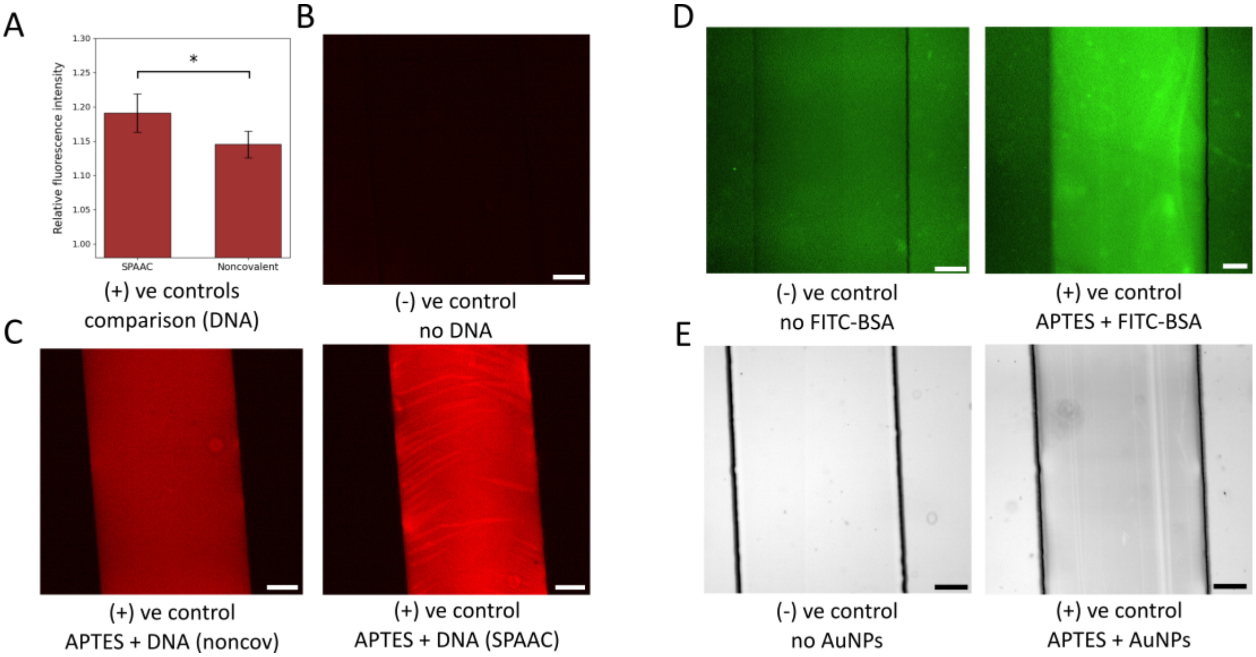


**Figure S7:** Supplementary positive and negative patterning controls on glass-PDMS devices. Negative controls were channels coated with PC PEG without adding cargos (DNA, FITC-BSA, gold nanoparticles (AuNPs); positive controls contained no PC PEG, so cargos could bind the entire channel. **(A)** Positive control comparison of SPAAC and noncovalently-bound azide-modified Cy3-DNA; fluorescence was normalised to background fluorescence outside channels (Mann-Whitney U-test, n=5). Representative images of **(B)** negative and **(C)** positive DNA patterning controls. **(D)** Representative images of negative and positive controls for noncovalent FITC-BSA patterning. The color scale is equivalent to Fig. 4E. **(D)** Representative images of negative and positive controls for noncovalent AuNP patterning, with 1% Tween20. Images scale is equivalent to Fig. 4I. Scale bars: 100 µm.

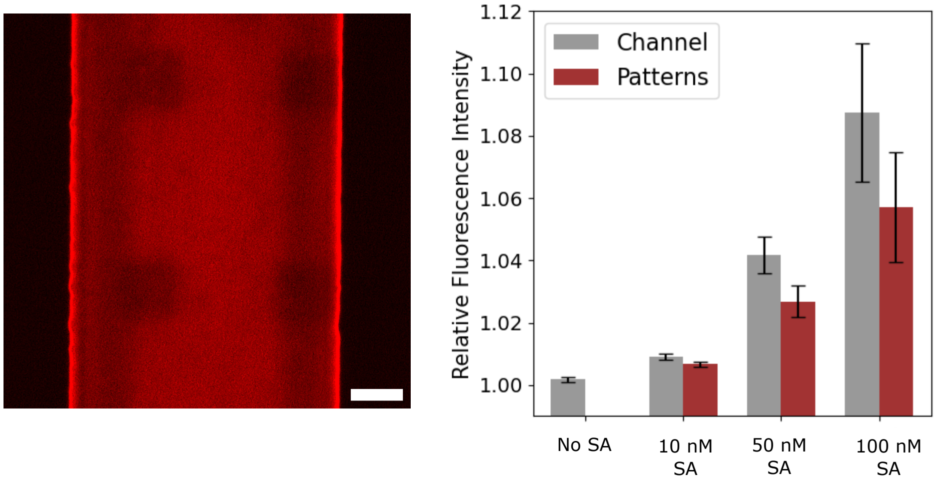


**Figure S8:** ATTO550-Streptavidin (SA) patterning. Devices were functionalized with PC PEG (with a DBCO head group), UV-patterned using a mask aligner, and ATTO550-SA loaded into devices at different concentrations. Devices were rinsed with PBS prior to imaging. Left: representative image of 100 nM ATTO550-SA patterns. We observed significant binding interactions outside patterns and on channel edges. Scale bar: 100 µm. Right: relative fluorescence intensity of patterns and channel (excluding patterns) produced with a range of ATTO550-SA concentrations (n=3). All concentrations led to 'negative' patterns, with higher nonspecific channel signal than pattern signal.

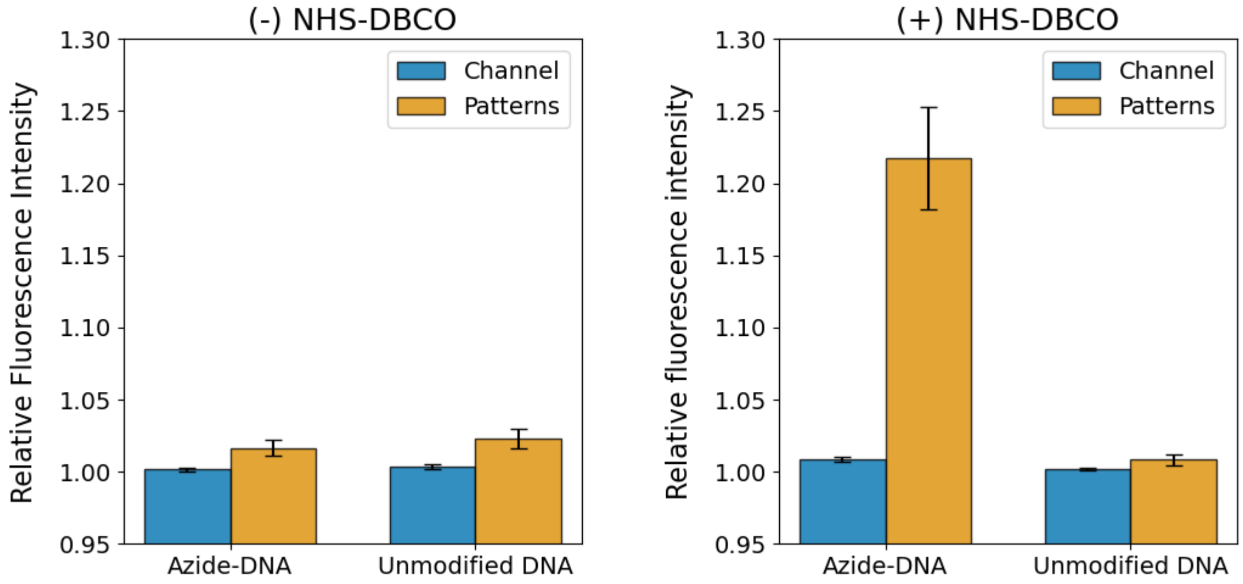


**Figure S9:** DNA channel and pattern quantification with and without NHS-DBCO. PC-PEG coated channels were UV-exposed, rinsed, and loaded with either NHS-DBCO or PBS buffer. After rinsing, the devices were incubated with either azide-modified DNA or unmodified DNA overnight, rinsed and imaged. Channel denotes unexposed areas, patterns denote UV-exposed areas. Bars represent mean and standard error (Left: n=15 (azide-DNA) and n=6 (unmodified DNA); right: n=28 (azide-DNA) and n=16 (unmodified DNA)).

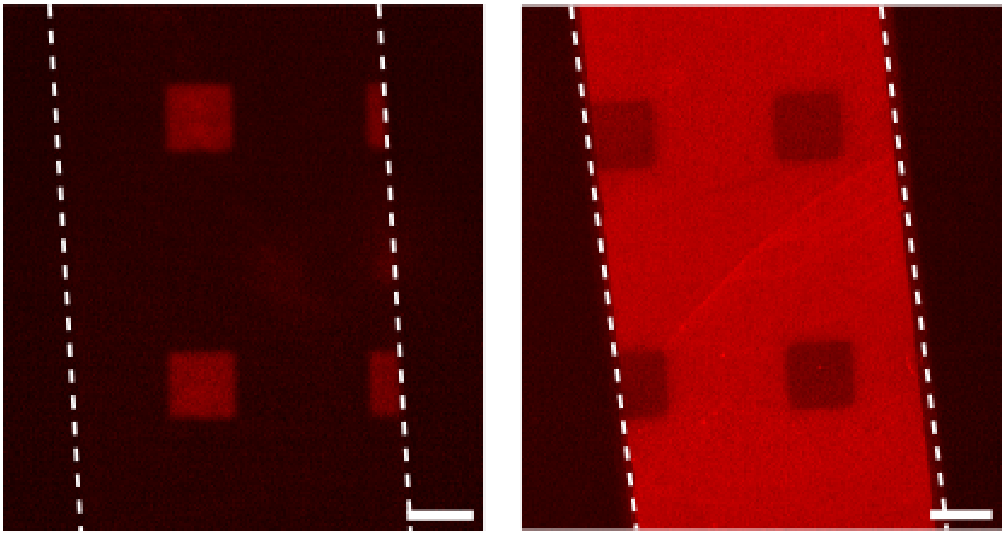


**Figure S10:** DNA positive and negative tone patterns with PC PEG containing a DBCO head group. Devices were first functionalized with PC PEG, patterned using a mask aligner to expose 50 $\mu$m squares, and then loaded with either unmodified DNA or azide-modified DNA. **Left:** unmodified DNA bound to UV-exposed areas through noncovalent interactions with APTES, forming 'positive' tone patterns. **Right:** Azide-modified DNA bound the DBCO-functionalized PC PEG via SPAAC, and more weakly bound the UV-exposed areas through noncovalent interactions with APTES. DNA was more densely bound to PC PEG than to APTES, leading to the formation of 'negative' tone patterns. Scale bar: 100 µm.

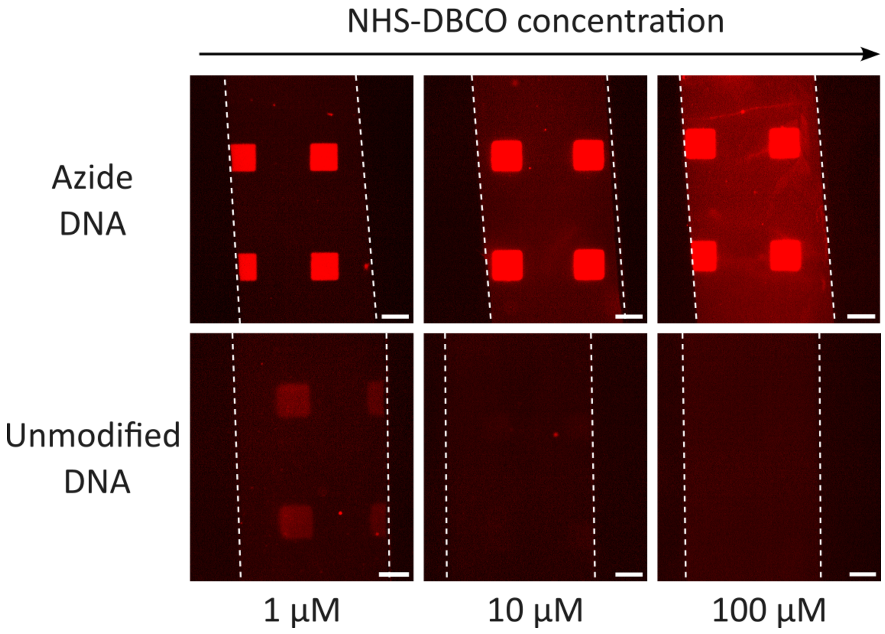


**Figure S11:** Effect of NHS-DBCO concentration on SPAAC and noncovalently bound DNA patterns. Devices were functionalized with PC PEG (functionalised with a biotin head group instead of DBCO to avoid cross-reactivity), UV-patterned with a mask aligner, and loaded with different concentrations of NHS-DBCO. Devices were subsequently rinsed with GnHCl and PBS, and incubated with either azide-modified or unmodified DNA. While higher NHS-DBCO concentrations more effectively blocked non-covalent DNA binding, they also resulted in higher off-target NHS-DBCO adsorption on unexposed surfaces. Scale bar: 100 µm.

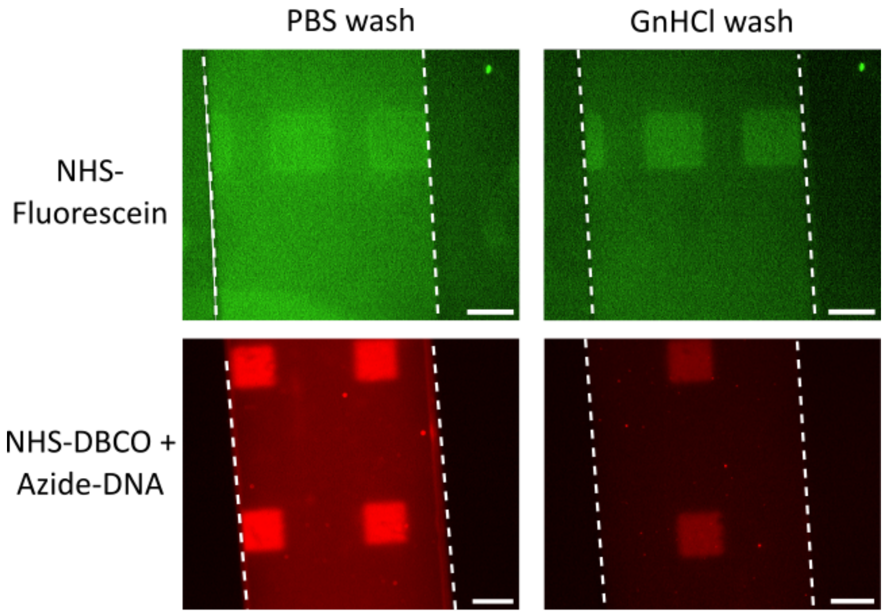


**Figure S12:** Effect of Gn-HCl rinsing on NHS-Fluorescein and NHS-DBCO. Devices were functionalized with PC PEG, UV-patterned with a mask aligner, and loaded with either NHS-Fluorescein or NHS-DBCO. After incubation, the devices were rinsed three times either with PBS or with 4 M GnHCl. Azide-modified DNA was then incubated overnight onto NHS-DBCO functionalized devices. Compared with PBS, Gn-HCl washing helped reduce nonspecific binding outside of UV-exposed patterns. Scale bar: 100 µm.

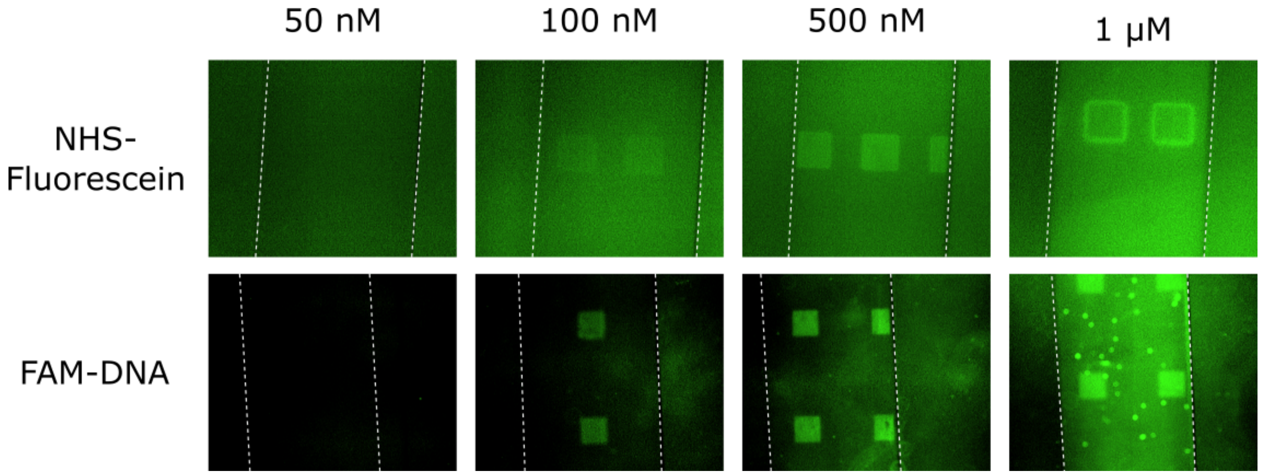


**Figure S13:** Effect of cargo concentrations on patterns. Devices were functionalized with PC PEG, UV-patterned with a mask aligner, and loaded with varying concentrations of NHS-Fluorescein or FAM-DNA, rinsed and imaged. At low cargo concentrations (50 nM), we were not able to detect fluorescence within intended patterns. At high cargo concentrations ($\geq 1\mu$M), we observed significant non-specific binding throughout the microchannels, both inside and outside intended patterns. Scale bar: 100 µm.

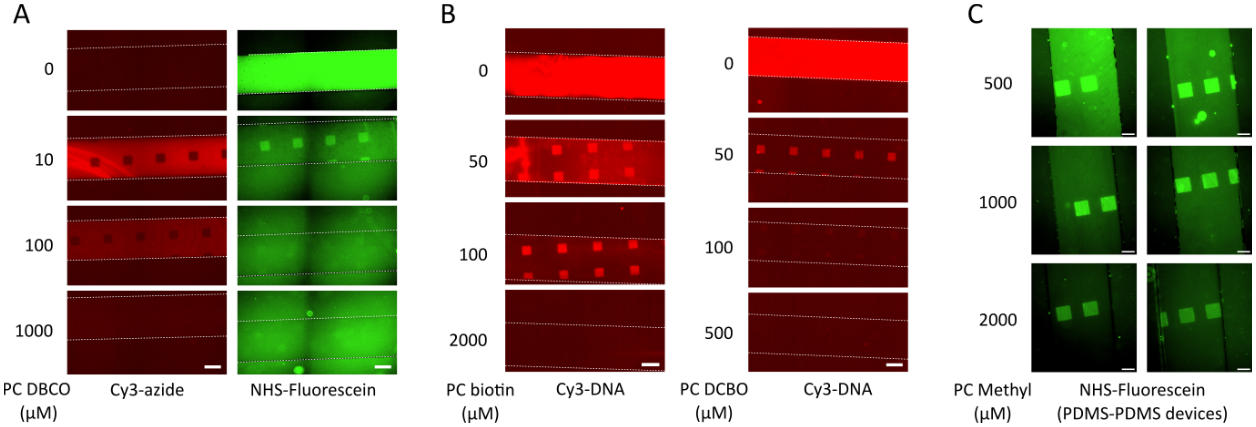


**Figure S14:** PC PEG concentration effects. Devices were functionalized with varying concentrations of PC PEGs, UV-patterned with a mask aligner, and functionalized using fluorescent probes. **(A)** PC PEG (containing a DBCO head group) concentration range functionalized with Cy3-azide (left) and NHS-Fluorescein (right). At high PC PEG DBCO concentrations ($\geq$ 100 $\mu$M), NHS-Fluorescein and cy3-azide no longer bound UV-exposed and unexposed areas, respectively. We hypothesised that at high PC PEG DBCO concentrations, complex layers formed on the surface through non-covalent interactions, leading to low accessibility of the DBCO group and obstructing photorelease of APTES amine groups in UV-exposed areas. **(B)** Comparison of PC PEG (biotin) and PC PEG (DBCO) concentration ranges on noncovalent DNA patterning. The optimal concentration differed between PC PEG compounds. **(C)** PC methyl concentration range on PDMS-PDMS devices, functionalized with NHS-Fluorescein). Concentration-dependant patterns also occurred on PDMS surfaces. Scale bar: 100 μm

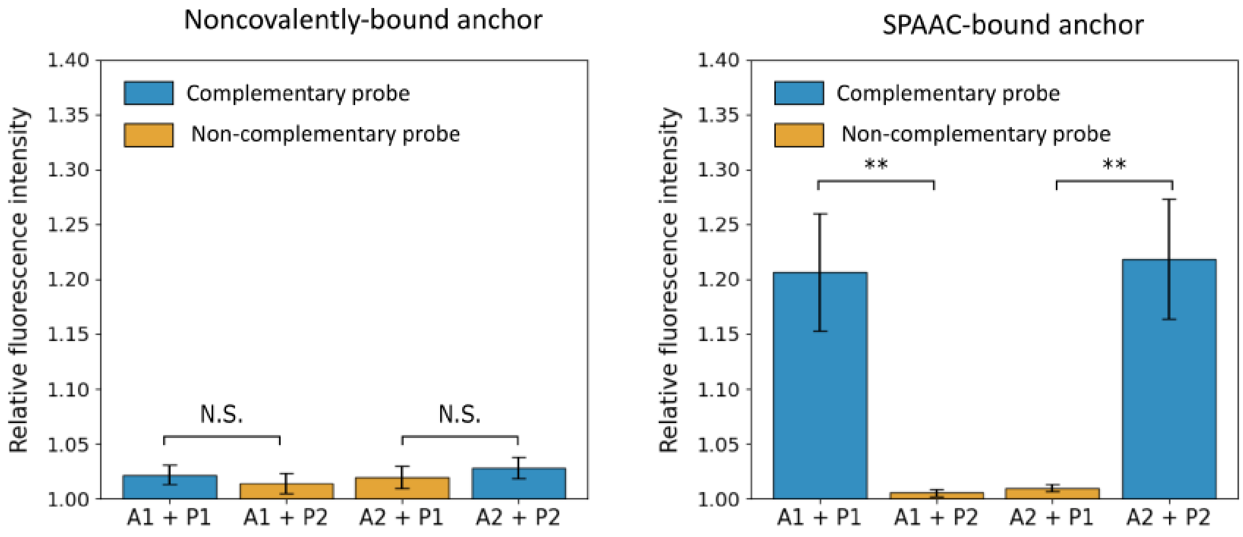


**Figure S15:** Quantification of pattern intensity for Cy3-DNA probe capture on noncovalently-bound (n=3-4) and SPAAC-bound DNA brushes (n=8-11) (corresponding to representative images in Fig. 5B). Only SPAAC brushes could reliably distinguish complementary and noncomplementary probes (Mann-Whitney U-test, $p<0.01$).

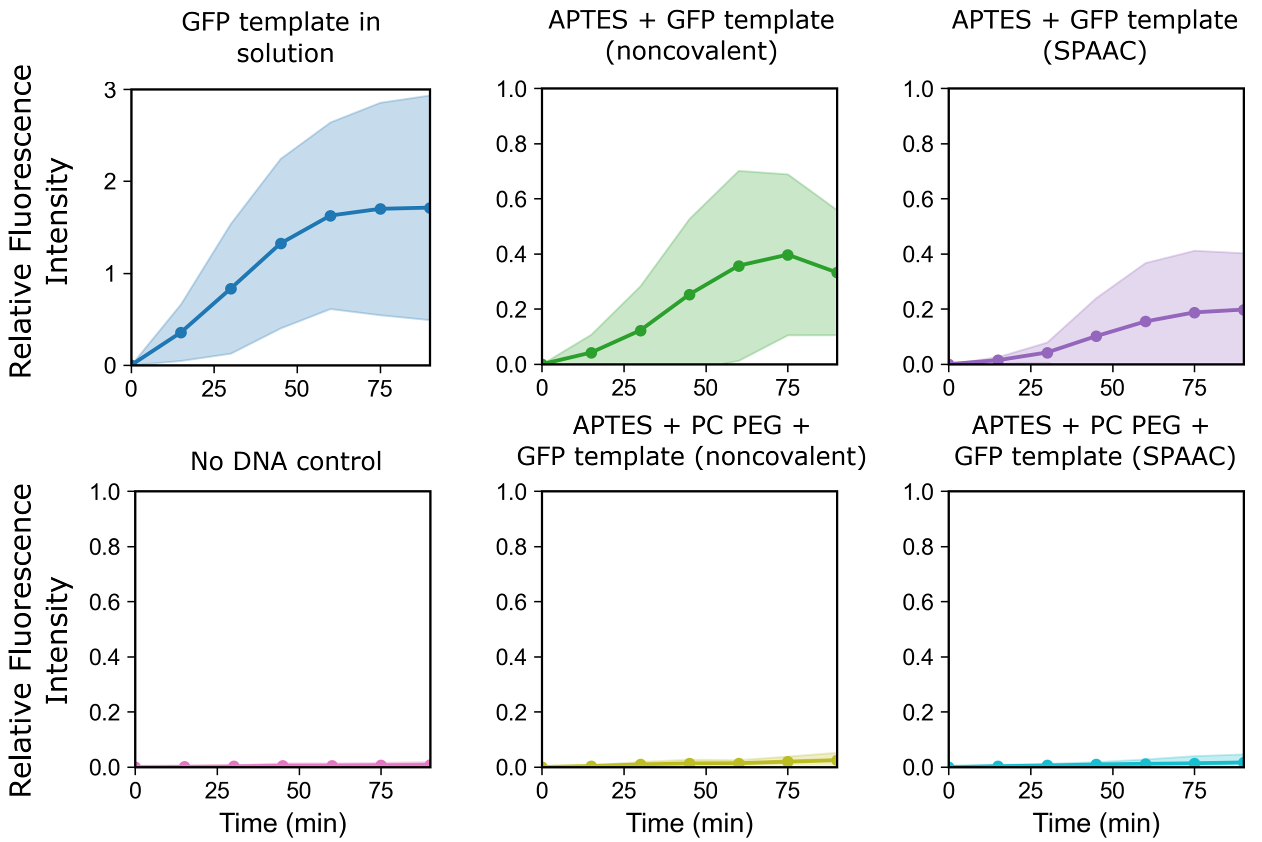


**Figure S16:** GFP expression trajectories. Microfluidic channels were coated with template DNA either with or without PC PEG, without UV patterning. PURExpress (cell-free gene expression mix) was then loaded into devices, and GFP signal monitored by microscopy. As a positive control, template DNA was pre-mixed with PURExpress before loading into channels (top left, n=3). As a negative control, PURExpress was loaded into channels without DNA (bottom left, n=7). We observed a modest increase in PURExpress autofluorescence over time. GFP expression from surface-bound DNA was first tested by immobilizing template DNA on APTES-coated devices via both noncovalent interactions and SPAAC anchoring (n=6). To test if PC PEG coating (without UV exposure) reduced DNA binding to the surface enough to reduce GFP expression, the same assays were carried out on devices coated with PC PEG before DNA immobilisation. In devices coated with PC PEG, we observed only a small increase in fluorescence that did not differ significantly from the gradual increase in PURExpress autofluorescence (n=8). Lines represent mean fluorescence intensity, shaded areas represent standard deviation. In each replicate, fluorescence from a manually defined area within the microchannel was averaged and normalised by dividing to the background fluorescence outside of the microchannel.

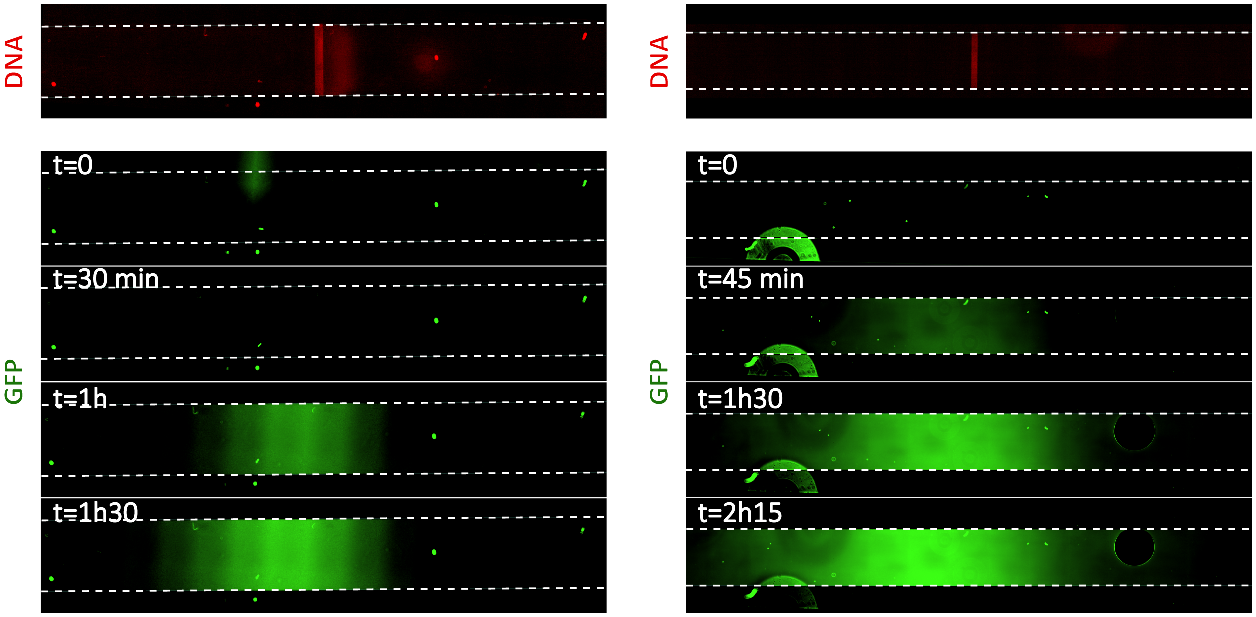


**Figure S17**: Replicates of GFP expression from patterned DNA, without fluorescent beads. A GFP template stripe was patterned in the centre of microfluidic channels (top). PURExpress when then loaded into microfluidic devices at t=0, and imaged over time (bottom). While peaks in GFP signal did emerge near the DNA stripe, a lateral shift was observed in all replicates.

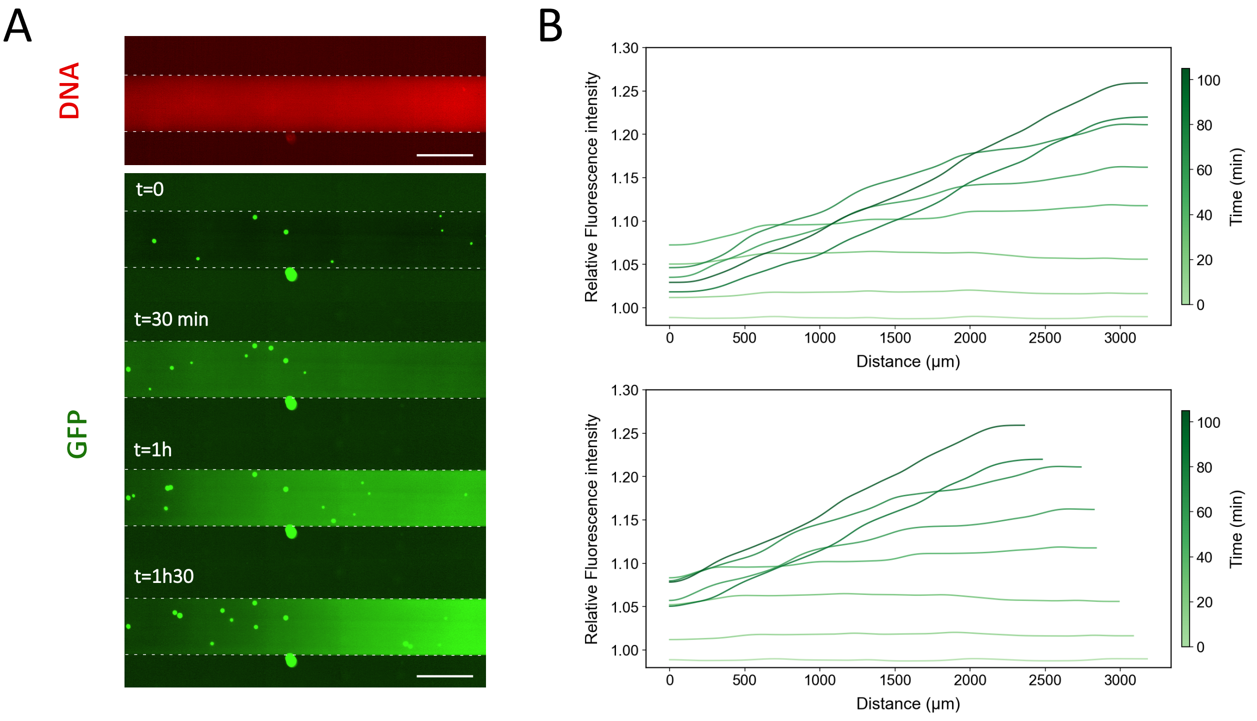


**Figure S18**: Representative timelapse of GFP expression from a uniformly-coated microfluidic channel. **(A)** A microfluidic channel was uniformly coated with the GFP template by incubating DNA directly on APTES, excluding the PC PEG patterning step. PURExpress (containing FAM beads to track fluid flow) was then loaded into devices and monitored over time. **(B)** GFP expression over time across the channel x-axis, with (bottom) and without (top) drift correction (calculated from bead movement). In uniformly-coated channels, we did not observe the peak-like GFP profile observed in patterned devices. A slope was observed after approximately 1h, perhaps due to a combination of fluid drift and PURExpress reagent depletion; DNA fluorescence intensity was slightly brighter on the right hand side of the image.

**Table S2: DNA sequences.**

| Name | Sequnce (5' - 3') | Modification |
|---|---|---|
| S1 | TTT GCG GAA TTG TAT GGT AGG TG | 5' Azide (optional) |
| S1' | TTT CAC CTA CCA TAC AAT TCC GC | 5' Cy3 |
| S2 | TTT ACT TAA CGA ACC GAG CTG TC | 5' Azide (optional) |
| S2' | TTT GAC AGC TCG GTT CGT TAA GT | 5' Cy3 |
| S3 | GAT GTG TCC AGT AGT CCG TA | 3' Azide (optional) |
| S3' | TAC GGA CTA CTG GAC ACA TC | 5' 6-FAM |
| F primer | TTT TCA ATA GGC TTG TGT CGA G | 5' ATTO550 |
| R primer | TTT TCA ATA GGC TTG TGT CGA G | 5' Azide (optional) |

**GFP template design:** *Design: spacer – T7 promoter – eGFP – T7 terminator – spacer*

**GFP Sequence:** TCAATAGGCTTGTGTCGAGCATTCAGGATCCCTTAATACGAC TCACTATAGGGAGAGTTTAAAGTCAGTAAAGAGGAGAAATAGACATGGTGAGC AAGGGCGAGGAGCTGTTCACCGGGGTGGTGCCCATCCTGGTCGAGCTGGACGG CGACGTAAACGGCCACAAGTTCAGCGTGTCCGGCGAGGGCGAGGGCGATGCCA CCTACGGCAAGCTGACCCTGAAGTTCATCTGCACCACCGGCAAGCTGCCCGTG CCCTGGCCCACCCTCGTGACCACCCTGACCTACGGCGTGCAGTGCTTCAGCCG CTACCCCGACCACATGAAGCAGCACGACTTCTTCAAGTCCGCCATGCCCGAAG GCTACGTCCAGGAGCGCACCATCTTCTTCAAGGACGACGGCAACTACAAGACC CGCGCCGAGGTGAAGTTCGAGGGCGACACCCTGGTGAACCGCATCGAGCTGAA GGGCATCGACTTCAAGGAGGACGGCAACATCCTGGGGCACAAGCTGGAGTACA ACTACAACAGCCACAACGTCTATATCATGGCCGACAAGCAGAAGAACGGCATC AAGGTGAACTTCAAGATCCGCCACAACATCGAGGACGGCAGCGTGCAGCTCGC CGACCACTACCAGCAGAACACCCCCATCGGCGACGGCCCCGTGCTGCTGCCCG ACAACCACTACCTGAGCACCCAGTCCGCCCTGAGCAAAGACCCCAACGAGAAG CGCGATCACATGGTCCTGCTGGAGTTCGTGACCGCCGCCGGGATCACTCTCGG CATGGACGAGCTGTACAAGTAAACCCGTGGTGGCGGCGGCAGCAGCGGTCCGC GTCCTCGTGGTACCCGCGGTAAGGGTCGTCGCATTCGTCGTTAATAATGGTAC CTACTAGCATAACCCCTTGGGGCCTCTAAACGGGTCTTGAGGGGTTTTTTGAC AGTCTGGAGTCGTAACAG
